# A genome-scale yeast library with inducible expression of individual genes

**DOI:** 10.1101/2020.12.30.424776

**Authors:** Yuko Arita, Griffin Kim, Zhijian Li, Helena Friesen, Gina Turco, Rebecca Y. Wang, Dale Climie, Matej Usaj, Manuel Hotz, Emily Stoops, Anastasia Baryshnikova, Charles Boone, David Botstein, Brenda J. Andrews, R. Scott McIsaac

**Affiliations:** Terrence Donnelly Centre for Cellular and Biomolecular Research, University of Toronto, Toronto, Ontario M5S 3E1, Canada; Department of Molecular Genetics, University of Toronto, Toronto, Ontario M5S 3E1, Canada; RIKEN Centre for Sustainable Resource Science, Wako, Saitama, Japan; Calico Life Sciences LLC, South San Francisco, CA 94080, USA

**Keywords:** BAR-seq, gene overexpression, yeast genomics, yeast mutant array, CRISPRi

## Abstract

The ability to switch a gene from off to on and monitor dynamic changes provides a powerful approach for probing gene function and elucidating causal regulatory relationships, including instances of feedback control. Here, we developed and characterized YETI (Yeast Estradiol strains with Titratable Induction), a collection in which 5,687 yeast genes are engineered for transcriptional inducibility with single-gene precision at their native loci and without plasmids. Each strain contains Synthetic Genetic Array (SGA) screening markers and a unique molecular barcode, enabling high-throughput yeast genetics. We characterized YETI using quantitative growth phenotyping and pooled BAR-seq screens, and we used a YETI allele to characterize the regulon of *ROF1,* showing that it is a transcriptional repressor. We observed that strains with inducible essential genes that have low native expression can often grow without inducer. Analysis of data from other eukaryotic and prokaryotic systems shows that low native expression is a critical variable that can bias promoter-perturbing screens, including CRISPRi. We engineered a second expression system, Z_3_EB42, that gives lower expression than Z_3_EV, a feature enabling both conditional activation and repression of lowly expressed essential genes that grow without inducer in the YETI library.

## Introduction

With its facile genetics and rapid division rate, the budding yeast *Saccharomyces cerevisiae* has been a leading model system for systematic studies of gene function [1]. To date, the most common genetic approach for exploring the biological roles of genes is to study phenotypes associated with loss of function alleles. The first genome-wide gene deletion strain collection was constructed in budding yeast, enabling a broad range of functional profiling studies [2]. Because ~20% of all genes are essential, causing lethality when deleted in a haploid cell [2], exploring loss-of-function of essential genes in this context requires the generation of conditional alleles. Conditional knockdown of gene function using degrons, transcriptional repression, and temperature sensitivity have been employed to investigate the role of essential genes, but each strategy has its own drawbacks, one of the most serious of which is general perturbations to cellular physiology associated with changes of environmental conditions [3–7].

Another systems-wide approach for studying gene function is gene overexpression, which can produce gain-of-function phenotypes and be used to study both essential and non-essential genes. Several gene overexpression plasmid collections with genes under the control of their endogenous promoters on high copy plasmids have been constructed [8,9]. Gene expression in these collections is not conditional, limiting phenotypic analysis and precluding the study of genes whose overexpression inhibits growth. To observe more dynamic overexpression phenotypes, the *GAL1* promoter, a strong inducible promoter that can be easily activated by the addition of galactose to glucose-free culture medium, has been used to construct a number of gene-overexpression plasmid collections [9–11]. However, *GAL1* promoter-based overexpression systems also have significant drawbacks, including the cell-to-cell variation of expression associated with replicative plasmids, non-graded induction, and the requirement of a metabolic signal for activation [12].

To address issues with yeast gene overexpression systems, we previously engineered a genome-integrated, conditional β-estradiol-inducible gene expression system in budding yeast [13–15]. In this system, an artificial transcription factor consisting of the modular zinc-finger DNA-binding domain, human Estrogen Receptor, and VP16 activation domain, is constitutively expressed. We refer to these artificial transcription factors as “ZEVs”. A synthetic promoter, which contains binding sites for ZEV variants, is inserted upstream of an open reading frame, displacing the endogenous promoter. The level of activity of the artificial transcription factor is controlled by β-estradiol concentration and enables regulated expression from the corresponding promoter. Since β-estradiol is not a yeast metabolite or signaling molecule, cellular metabolism is not perturbed [13]. ZEVs have been widely used for basic and applied research, including studies of gene regulatory networks [16–18], individual gene function [19–25], gene regulation [26–30], metabolic engineering [31], synthetic biology [32–39], biocontainment [40], living materials [41], high-throughput screening [42,43], and they have also been adapted to fission yeast [44–46] and *Pichia pastoris* [47].

ZEVs allow the rapid induction of a single gene in any environment, which provides a system for tracking how induction of gene expression is directly linked to a cellular response, something that cannot be achieved with deletion mutants [16,17,48]. To generate a systems-level reagent set for molecular and cellular analysis using the ZEV system, we constructed the YETI collection, in which nearly every gene in budding yeast is inducible with Z_3_EV, a ZEV variant that utilizes the 3-finger, Zif268 zinc finger from mouse to bind DNA. We thoroughly characterize these strains and provide details for researchers interested in using this collection. In total, we integrated a uniquely-barcoded β-estradiol-regulated promoter in front of 4,665 non-essential genes and 1,022 essential genes in a heterozygous diploid background, and recovered 5,670 Z_3_EV-driven genes in a haploid background using the synthetic genetic array (SGA) selection system [49]. By combining this collection with automated yeast genetics and dynamic growth profiling, we identified 985 genes whose overproduction reduces cell fitness at higher levels of β-estradiol. Additionally, we identified 46 genes whose expression levels affect fitness in a non-monotonic fashion, demonstrating the utility of this collection for genome-scale exploration of fitness landscapes. While more than half of strains with Z_3_EV-driven essential genes were not able to grow as haploids in the absence of β-estradiol, another subset was viable even without inducer. These findings motivated us to develop a second expression system – the Z_3_EB42 system – which involved re-engineering Z_3_EV as well as its target promoter to generate a gene regulation system that gives lower expression and is more extensively repressed in the absence of inducer. The YETI strain collection provides a new and comprehensive platform for interrogating yeast gene function and dynamics.

## Results

### A genome-scale collection of inducible alleles

To construct a genome-scale collection of strains expressing β-estradiol-inducible alleles, we first constructed a parental strain expressing the Z_3_EV transcription factor. Our diploid parental strain, Y14789, was based on the RCY1972 strain. We chose RCY1972 as it is deleted for the *HIS3* locus, making it compatible with SGA methodology, but otherwise prototrophic, enabling studies of yeast cell growth and other phenotypes in a variety of conditions [50]. The strain also carries a wild-type *HAP1* gene, which encodes a transcription factor that localizes to both the mitochondria and nucleus and is required for regulation of genes involved in respiration and the response to oxygen levels [51]. Strains derived from S288C carry a Ty1 element insertion in the 3’ region of the *HAP1* coding sequence, creating a *HAP1* allele that acts as a null for cytochrome *c* expression and leads to mitochondrial genome instability [51]. Previous work has shown that removal of the Ty element, which repairs the *HAP1* gene, increases sporulation efficiency dramatically [52]. To select for strains that carried a wild-type *HAP1* gene, the gene encoding the Z_3_EV transcription factor was integrated next to the repaired *HAP1* together with a *natMX* selectable marker in Y14789. Each strain also carries the SGA marker system (*can1*Δ::*STE2pr-Sphis5* and *lyp1*Δ), enabling automated, array-based genetics. Finally, we engineered a DNA template on which the *URA3* gene is linked to Z_3_pr for creating genomically integrated promoter fusions. The components of the β-estradiol gene expression system are outlined in **Figure 1A**.

**Figure 1:**
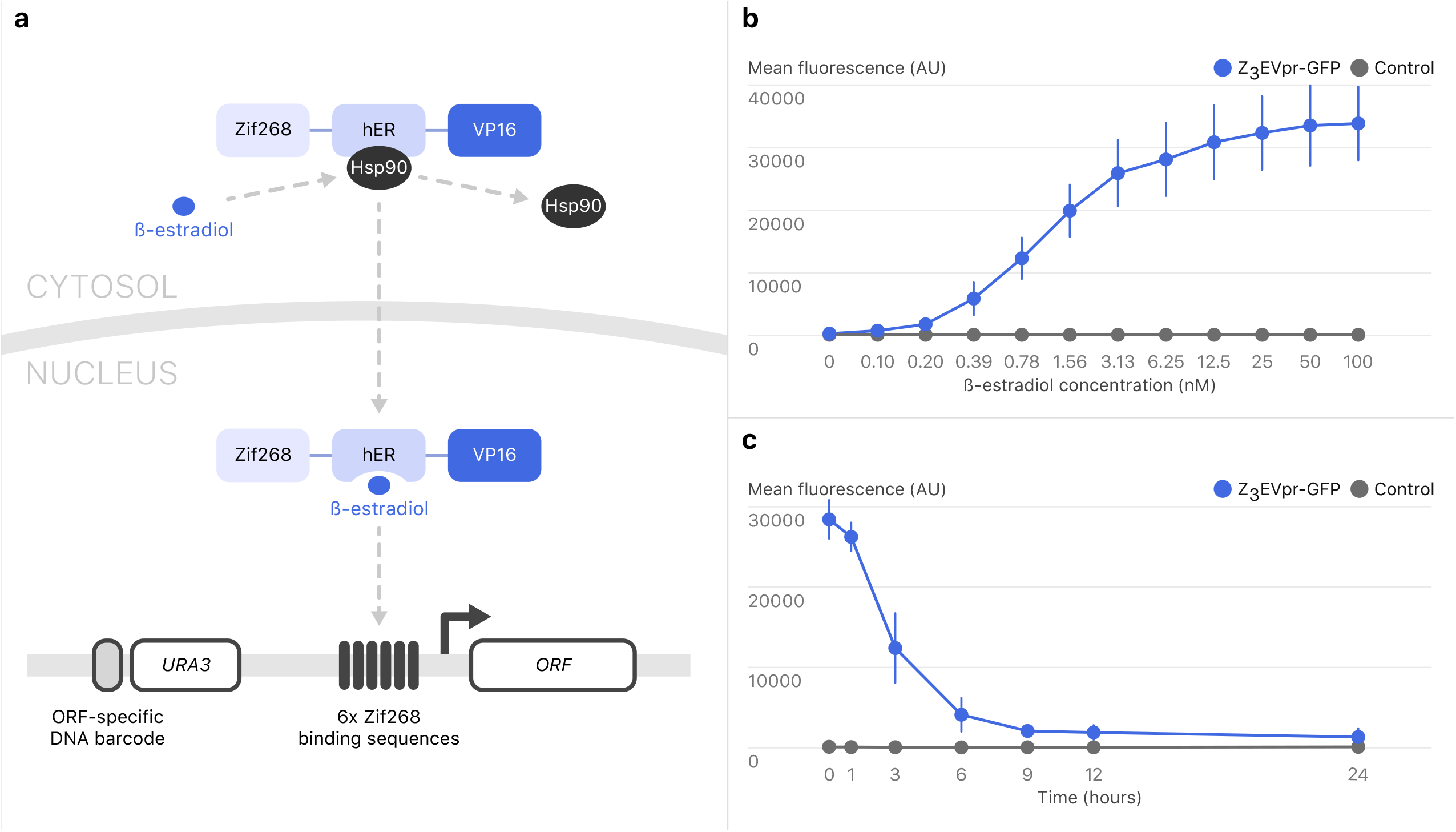
The Z_3_EV system. **A.** Outline of the β-estradiol-inducible gene expression system. Z_3_EV is composed of a 3-zinc-finger DNA-binding domain (Zif268), human estrogen receptor domain (hER) and transcription activation domain (VP16). β-estradiol displaces Hsp90 from the estrogen receptor, allowing Z_3_EV to translocate to the nucleus and induce gene expression. Zif268 binds preferentially to a sequence that is present in six copies in Z_3_pr. In the strain collection, gene-specific DNA barcodes are flanked by universal primer sequences: 5’-GCACCAGGAACCATATA-3’ and 5’-GATCCGCTCGCACCG-3’. **B.** GFP intensity as a function of β-estradiol concentration. Strains with an integrated Z_3_pr driving GFP (Y15292; blue) and a control strain (Y15477; grey) were incubated with a concentration series of β-estradiol in YNB for 18 hours and then cells were fixed. GFP intensity was measured by flow cytometry. Error bars represent standard deviation for three replicates. **C.** Y15292 (blue) and Y15477 (grey) cultures were induced with 10 nM β-estradiol for 18 hours. Cells were washed and β-estradiol was removed from the medium at time = 0 hours. Error bars represent standard deviation for three replicates.

To test the β-estradiol concentration- and time-dependent expression of a gene regulated by the Z_3_EV transcription factor in our strain background, we first inserted the Z_3_pr in front of *GFP* and measured GFP fluorescence intensity by flow cytometry. Expression of the Z_3_pr-*GFP* reporter gene increased in a concentration- and time-dependent manner (**Figure 1B, Figure S1A**). Following removal of β-estradiol, the GFP signal decreased by ~50% within three hours and decreased to near-background levels within 24 hours, demonstrating that expression of the GFP reporter gene was dependent on β-estradiol concentration and treatment time (**Figure 1C**). We also tested whether mutant phenotypes could be complemented by cognate genes expressed from the Z_3_ promoter. We engineered strains in which *TPS2* or *LEU2* were placed downstream of Z_3_pr and in the absence of β-estradiol, the resultant strains displayed known phenotypes associated with *leu2Δ* (**Figure S1B**) and *tps2Δ* (**Figure S1C**), leucine auxotrophy [53] and heat sensitivity [54], respectively, but grew equivalently to WT cells in the presence of β-estradiol.

Following these successful characterizations of our constructs, we engineered a genome-wide collection in which the endogenous promoters of individual genes were replaced by inserting the Z_3_pr just upstream of the start codon of each ORF in a heterozygous diploid strain carrying the Z_3_EV transcription factor under the control of the constitutive *ACT1* promoter, which we linked to a *natMX* marker. Importantly, the *URA3* gene marking the Z_3_ promoter in each strain is linked to a unique DNA molecular barcode such that the resulting genome-wide β-estradiol-inducible strain collection is compatible with pooled screening approaches. Promoter insertions were confirmed by PCR, and further quality control with a subset of specific strains was carried out using whole-genome sequencing (see Supplement, <u>**Table S1**</u>). In total, we constructed 1,022 diploid strains carrying β-estradiol-inducible alleles of essential genes (corresponding to 97.1% of essential genes) (<u>**Table S2A**</u>) and 4,665 diploid strains expressing β-estradiol-inducible alleles of non-essential genes (corresponding to 98% of non-essential genes) (<u>**Table S2B**</u>).

For the non-essential genes, we recovered haploid derivatives by sporulating the heterozygous diploid strains and germinating haploid meiotic progeny on SGA selection medium (**Table S2C**). We refer to these haploid strains as the YETI non-essential (YETI-NE) panel. The set of strains with essential genes under the control of the β-estradiol-inducible promoter are maintained as diploids. We refer to these strains as the YETI essential (YETI-E) panel. We were unable to construct 127 strains (30 YETI-E diploids, 87 YETI-NE diploids, and 10 YETI-NE haploids; <u>**Table S2D**</u>). We estimate the percentage of defective strains to be <5% (see Methods for details).

### Quantifying the relationship between inducer level and transcript level

A useful feature of the Z_3_EV system is the potential to either knock down or induce a gene of interest depending on the level of β-estradiol inducer, which enables comprehensive analysis of the relationship between gene expression and phenotype by finely tuning transcription. To assess this property of the collection, we selected Z_3_pr alleles of 18 non-essential genes with a range of native expression levels (from 3 to 2173 TPM [transcripts per million]). Following growth in the presence of various concentrations of β-estradiol, we quantified mRNA expression levels of the β-estradiol-regulated genes using RNA-seq. All genes had qualitatively similar responses to β-estradiol concentration: very low expression at β-estradiol concentrations from 0-1 nM, and then an increase in expression between 4 and 16 nM, followed by a plateau at β-estradiol concentrations beginning at 16-64 nM (**Figure S2A**). However, the actual level of transcript produced at each β-estradiol concentration varied. For this gene panel, maximum expression levels ranged from ~30-40 TPM for some genes (e.g., *ATG4*, *SNT1* and *SRS2*) to over 1000 TPM for others (e.g., *ASC1*, *BAT1*, *THR1* and *VMA3*) at >16 nM β-estradiol. Peak expression levels were correlated with native transcript levels (Pearson correlation = 0.74), which is reminiscent of the gene-dependent expression variation reported with a GALpr plasmid collection [55]. For genes in the panel whose native expression was <250 TPM, the maximum expression output from Z_3_EV and the level of native expression followed a simple linear relationship (**Figure S2B**). Linearity broke down for the two highly expressed genes we tested, *ASC1* and *VMA3*; even at saturating concentrations of β-estradiol, Z_3_EV-induced expression was lower than or equal to native expression levels. We conclude that Z_3_EV can be used for titrating gene expression, but the exact number of transcripts produced depends on the target gene open reading frame and its genomic context.

### Growth patterns associated with β-estradiol-dependent regulation of essential genes

To begin our characterization of the YETI strain collection, we wanted to describe the possible behaviors of the β-estradiol-regulated promoter alleles. We first explored the growth characteristics of the essential gene alleles (YETI-E) using high resolution time-lapse imaging (see Methods for details) at twelve β-estradiol concentrations since, unlike non-essential genes, alterations in essential gene expression are expected to lead to an easily assayed growth phenotype (**Figure 2**). YETI-E strains were grown as heterozygous diploids, sporulated and haploid YETI-E strains were selected on medium with various β-estradiol concentrations. Growth profiles were then hierarchically clustered using a Chebyshev distance metric, which revealed five distinct clusters or promoter behaviors (**Figure 2, <u>Table S3A</u>**). The largest cluster contained 49% of YETI-E strains, each of which showed a dosage-dependent growth response, with no growth in the absence of inducer, and improved growth with increasing β-estradiol concentrations (Cluster 5, **Figure 2**). A second smaller cluster showed a similar initial behavior, with growth depending on the presence and concentration of inducer, but with eventual growth inhibition, consistent with dosage toxicity (Cluster 4 with 4.7% of strains). Thus, more than half (53.7%) of the YETI-E strains exhibited the expected β-estradiol-dependent gene expression that is clearly ‘tunable’ by inducer concentration.

**Figure 2:**
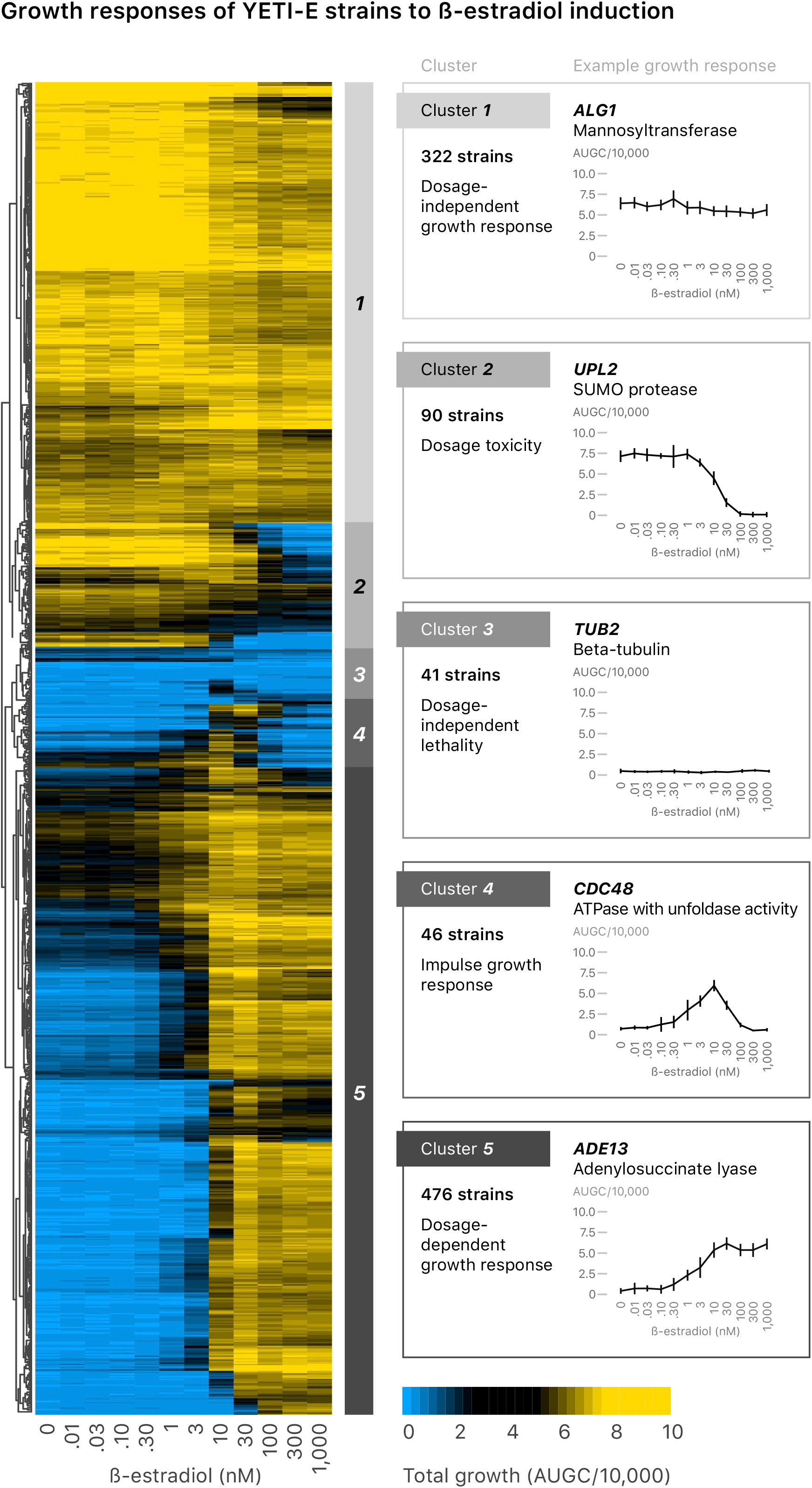
Hierarchical clustering of YETI-E growth patterns. The YETI-E strains were grown on SC SGA selection medium with monosodium glutamate as a nitrogen source at 12 β-estradiol concentrations (0, 0.01, 0.03, 0.1, 0.3, 1, 3, 10, 30, 100, 300, 1000 nM) and clustered by growth profile. Growth is measured as the area under the growth curve (AUGC); blue represents little growth and yellow represents more growth. Strains were clustered using a Chebyshev distance metric, resulting in 5 clusters (numbered 1-5). Representative strains for each of the five clusters and their growth patterns are plotted to the right of the clustergram.

Most remaining strains in the YETI-E collection had a dosage-independent growth response, including a large set of strains (33%) that grew in the absence of β-estradiol and showed no response to increased β-estradiol concentration (Cluster 1 – dosage-independent growth), and a smaller set (9.2%) that grew in the absence of inducer, with growth inhibition at higher concentrations (Cluster 2 – dosage toxicity). Finally, a small group of strains (4.2%), with no obvious functional features in common, failed to grow well regardless of the inducer concentration, suggesting these strains have a non-functional promoter (Cluster 3 – dosage independent lethality) (**<u>Table S3B</u>**).

Roughly half of YETI-E strains grew in the absence of β-estradiol, indicating sufficient basal gene expression to support essential gene function (**Figure 2**, **Table S3A**). Our transcriptome analysis of a panel of 18 Z_3_pr-driven alleles showed that transcript levels were reduced to 30%, on average, of their native levels in the absence of inducer (**Figure S2A, Figure S3**), and similar results were found with 201 β-estradiol-controlled transcription factors [16]. The ability of low levels of YETI-E allele expression in the absence of inducer to support growth may correlate with the levels of native gene expression. To test this possibility, we divided essential genes into six bins based on native gene expression levels [56] and plotted the growth distribution of each bin. Genes with low transcript levels had significantly better growth in the absence of β-estradiol than genes with high transcript levels (KS test, p < 1.9 × 10^−20^; **Figure 3A**, **Table S4A**). The converse is also true: YETI-E strains that grew poorly at 0 nM β-estradiol contain regulated alleles of genes that were more highly expressed from their native promoters than strains that grew well (**Figure S4A**). We conclude that though Z_3_pr is extremely weak in the absence of β-estradiol, it still promotes enough transcription of many essential genes that are normally lowly expressed to facilitate growth.

**Figure 3:**
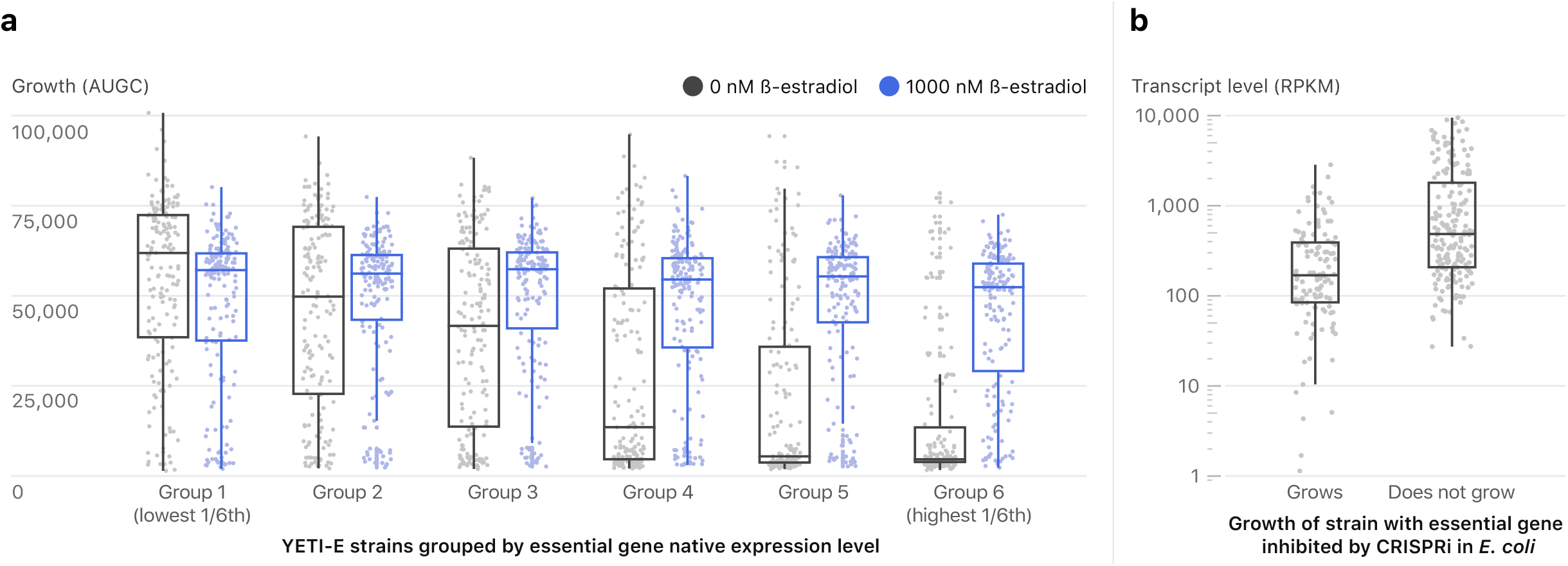
Synthetic control of growth depends on native gene expression levels. **A.** Yeast strains in which Z_3_EV targets essential genes are more likely to grow in the absence of inducer if the Z_3_EV-controlled gene is lowly expressed. The boxplots show YETI-E AUGC (area under the growth curve) values from 0 nM (blue) and 1000 nM (grey) β-estradiol experiments. The 6 bins (bins 1-6) are based on gene expression level of native genes [56]. **B.***E. coli* strains in which CRISPRi targets essential genes are more likely to grow if the targeted essential gene is lowly expressed. Boxplots show the distribution of gene expression levels for *E.coli* genes tested in a CRISPRi screening experiment [59]. The genes are grouped into essential genes whose repression by CRISPRi inhibits growth (does not grow) or fails to inhibit growth (grows). Gene expression data from Gene Expression Omnibus (GSE67218). RPKM median values are 166 and 503, for the “Grows” and “Does not grow” classes, respectively.

We compared our results to the growth characteristics and gene expression patterns of strains in other yeast collections that enable conditional induction or repression of essential genes. In the Tet-off collection [3], the native promoter of a target gene is replaced by a constitutive promoter that contains elements that allow it to be repressed in the presence of doxycycline. Of the 453 strains tested in both systems, 224 had strongly reduced growth in the presence of doxycycline (dox-responsive) and 229 were either not affected or were minimally affected by doxycycline (dox-non-responsive). We found that 74% of dox-non-responsive genes were also β-estradiol-non-responsive (union of Clusters 1+2, **Figure 2**). In contrast, only 30% of dox-responsive genes were β-estradiol-non-responsive, a ~1.6-fold reduction from what is expected by random chance (**Table S4B**). Dox-responsive genes were also more highly expressed than dox-non-responsive genes (p-value = 0.002, two-sided t-test, **Figure S4B**). Thus, native expression can influence one’s ability to achieve conditional growth with systems that both “turn down” (Tet) and “turn up” (Z_3_EV) gene expression.

We used the literature to explore if yeast strains bearing essential genes whose expression was perturbed in other ways had the same inverse correlation between native transcript level and fitness. The DAmP (Decreased Abundance by mRNA Perturbation) collection contains 842 essential genes whose 3′ untranslated region (UTR) is disrupted with an antibiotic resistance cassette, which can destabilize the corresponding transcript [57]. Fitness of the DAmP strains was assessed using a competitive growth assay. Consistent with our findings with the Z_3_EV and Tet-off systems, strains containing DAmP alleles that had closer-to-wild-type fitness tended to be in genes with lower native expression levels than DAmP strains with reduced fitness (**Figure S4C**; **Table S4C**, p-value = 0.002, two-sided t-test). N-terminal heat-inducible-degrons can also be used to target a fusion protein for degradation at 37°C in a Ubr1-dependent manner [7]. Of 94 degron-tagged essential genes, those for which the degron had no effect on temperature-sensitivity had lower native expression than those for which the degron caused a temperature-sensitive phenotype (**Figure S4D**; **Table S4D**, p-value = 0.0001, two-sided t-test). Finally, we looked at the effects of a new inducible CRISPR-interference (CRISPRi) library on growth of strains with essential genes [58]. CRISPRi depletion scores for highly-expressed essential genes were significantly stronger (more negative) than those of lowly-expressed essential genes (**Figure S4E**; **Table S4E**, p-value = 3.6 × 10^−5^, two-sided t-test).

To test if native expression level is also an important factor for achieving conditional growth for essential genes in organisms other than yeast, we investigated a recent CRISPRi pooled screen from *Escherichia coli* [59]. A useful feature of *E. coli* for this analysis is the availability of a deletion mutant collection from which a core set of essential genes has been determined [59,60]. In the CRISPRi screen, 62% of strains with essential genes scored as “non-dividing” (Class ND), i.e. these genes were inhibited enough by CRISPRi to block cell division. Thirty-eight percent of essential genes were not inhibited enough by CRISPRi to prevent cells from dividing (Class D). We find that Class ND genes were, on average, 5x more highly expressed than Class D genes (**Figure 3B; Table S4F**).

Collectively, we find that perturbations of expression of essential genes in eukaryotes and prokaryotes, including promoter replacement (Z_3_EV and Tet-off), 3’-UTR disruption, N-terminal degron and CRISPRi, do not have equal effects on fitness for all genes. Reducing expression of essential genes with high native transcript levels tends to cause fitness defects, while genes with low expression are more resistant to the effects of synthetic perturbation.

### Growth profiles associated with β-estradiol-dependent regulation of non-essential genes (YETI-NE)

By definition, failure to express a non-essential gene in a strain grown in a rich medium is not expected to produce a dramatic growth phenotype. However, some non-essential genes are required for growth in certain conditions, including metabolic auxotrophs. We defined metabolic auxotrophs as a set of 80 genes annotated in the *Saccharomyces Genome Database* (SGD) [61] that typically function in the biosynthesis of amino acids or nucleotides and whose deletion causes amino acid or nucleotide auxotrophy. Of these, 14 metabolic auxotrophs are not present in the YETI-NE collection and 24 metabolic auxotrophs are in pathways that are precluded from our analysis because they are part of the SGA selection protocol used in the construction of the YETI-NE collection (uracil, lysine, arginine, and histidine). Indeed, whole-genome sequencing of a subset of these SGA selection-pathway strains, grown on minimal medium under a condition where they should not actually propagate, revealed chromosome duplication or partial duplication events in 10 of the 17 strains (~59%) (**Table S1**), with most events (8/10) including the gene of interest, consistent with strong selection pressure resulting in aneuploidy [62]. For haploid YETI-NE strains outside of selection pathways that we sequenced, 4% (1 strain out of 23) had detectable aneuploidy. In total, 42 metabolic auxotrophs in the YETI-NE collection could be assessed for β-estradiol-dependent growth in auxotrophic conditions.

To explore the behavior of the auxotrophic allele panel in the YETI-NE collection, and to reveal any additional auxotrophies, we quantified the growth of the entire YETI-NE haploid panel on minimal YNB (yeast nitrogen base without amino acids or bases) and richer SC (yeast nitrogen base with all amino acids and bases added; see Supplement) solid medium in the absence or presence of β-estradiol (**Table S5A**). To quantify strain growth, we defined a metric called the Aux-score, which provides a measure of whether a non-essential gene behaves like an auxotroph on YNB medium: Aux-score_i_ = (G_*iM*1_/G_*iM*0_)/(G_*iR*1_/G_*iR*0_). G, *i*, *M*, *R*, 0, and 1 denote total growth, genotype, YNB, SC, 0 nM β-estradiol, and 1 nM β-estradiol, respectively (Aux-scores for all YETI-NE genes are shown in **Table S5B**). Z_3_pr-controlled auxotrophs should see a more dramatic growth improvement when β-estradiol is added to YNB than to SC, and thus, these strains should have large Aux-scores. Strains that have reduced growth on YNB medium in the presence of β-estradiol should have small Aux-scores (Aux-scores for YETI-NE genes are provided in **Table S5B**).

When Aux-scores were ranked from large to small values, as expected, most strains showed no difference in behaviour in the presence of β-estradiol on SC vs. YNB medium (Aux-scores of 1, **Figure 4**). However, clear outliers emerged, and thirty-eight of the 42-member metabolic auxotroph panel had Aux-scores greater than 1, indicating proper regulation by the ZEV promoter (the 24 top hits from the metabolic auxotroph panel are labeled in blue in **Figure 4**). Some strains with high Aux-scores were not known auxotrophs, and the list includes strains with ZEV alleles driving a variety of metabolic enzymes, as well as several components of the SWI/SNF chromatin modifying complex (the top 17 hits that are not part of the metabolic auxotroph panel are labelled in gray in **Figure 4**). We validated the growth profile of a strain containing a Z_3_pr allele of *SHM2,* which encodes a cytosolic serine hydroxymethyltransferase (**Figure S5A**). A series of growth experiments revealed that deletion of *SHM2* resulted in adenine auxotrophy, explaining why the Z_3_pr-*SHM2* strain required β-estradiol to grow on YNB, but not in adenine-containing SC medium (**Figure S5B, Figure S5C**). These examples illustrate that the subset of YETI-NE strains with phenotypes on YNB medium show β-estradiol dependent growth characteristics reflective of their functional roles.

**Figure 4:**
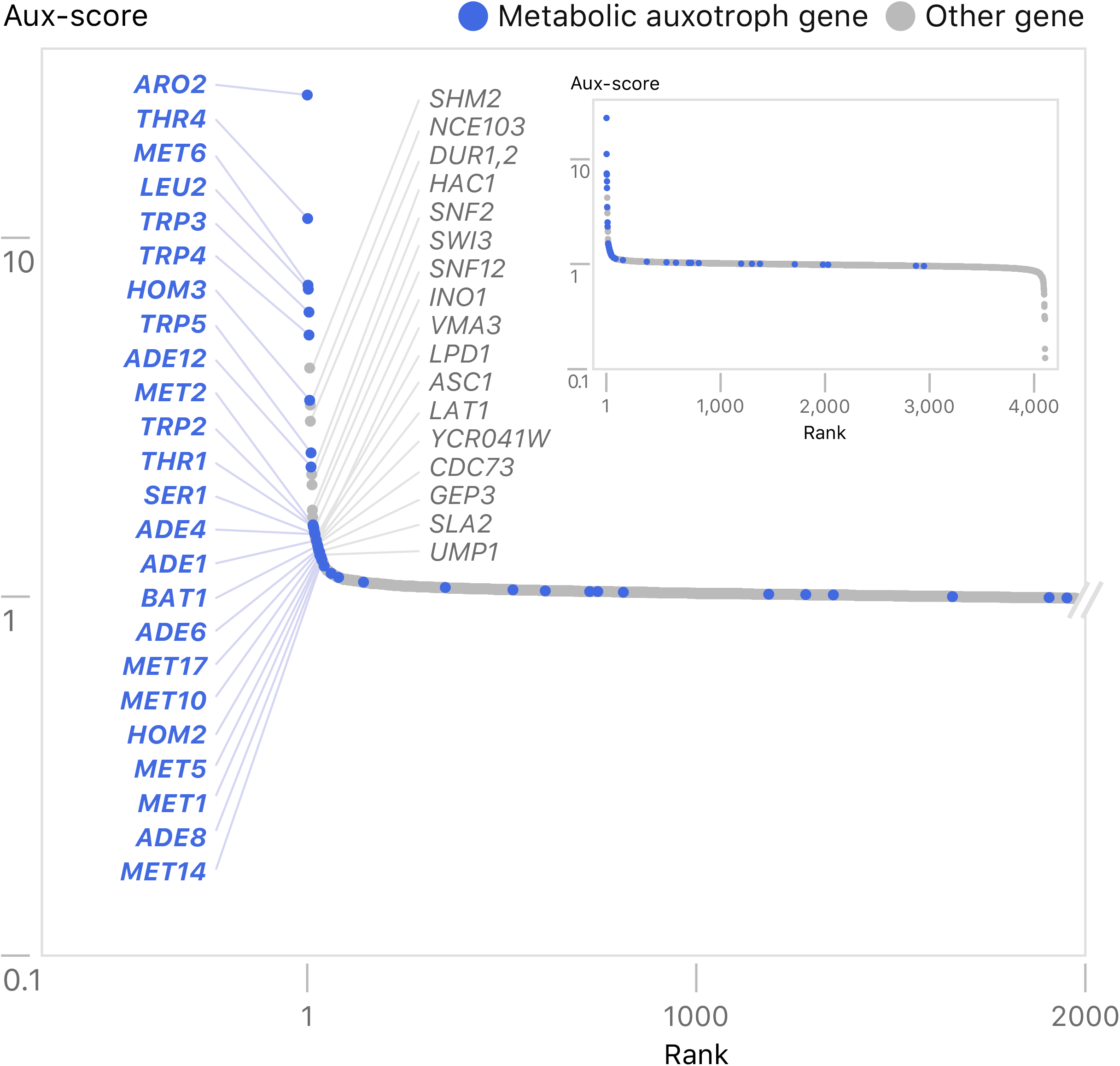
Comparing β-estradiol-dependent growth of strains expressing YETI-NE genes on YNB and SC. Distribution of Aux scores of YETI-NE strains. The y-axis shows the Aux score value (see main text and methods) measured for each YETI-NE strain displayed on the x-axis. YETI-NE strains are shown as grey dots with metabolic auxotroph panel members labeled in blue. The identities of the 41 YETI-NE strains with the highest Aux scores are listed. The inset shows the distribution pattern for all strains tested.

### Identification of genes that impair growth when overexpressed on plates

The Z_3_EV allele collection enables tunable regulation of gene expression and a systematic survey of the phenotypic consequences of gene over-expression. Aggregating plate-based growth data from the YETI-E and YETI-NE panels, we observed that ~17% (987/5671) of strains tested (specifically, 301 YETI-E stains and 686 YETI-NE strains) had reduced growth on SC medium at 100 nM β-estradiol as compared to growth at 0 nM β-estradiol (**<u>Table S6A</u>**, **Figure S6A-D**, see Methods). Amongst these strains, we saw a direct correlation between increased β-estradiol concentration and the severity of the growth defect. Previous studies of gene overexpression toxicity have explored various mechanisms of sensitivity to protein overproduction, including the balance hypothesis, which posits that deviations in the stoichiometry of protein complex members affect the complex’s overall function [63]. While conflicting results have emerged from different gene overexpression studies [10,64,65], the 987 toxic genes we identified show no enrichment (χ^2^ test, p = 0.6) for members of protein complexes (data from [66,67]), in agreement with the results of Sopko et al. [10]. Our list of toxic genes was enriched for genes with annotated roles in regulation of the mitotic cell cycle, DNA replication, microtubule cytoskeleton organization and chromosome segregation, all bioprocesses related to cell growth that have been reported previously in studies of overexpression toxicity (**Table S6B**, [10,11,55,65]). We also found enrichment of several GO function and component terms: transcription factors, nucleocytoplasmic carrier, SUMO transferase activity, and genes involved in P-body formation.

We examined whether physical properties of proteins were associated with overexpression toxicity. Intrinsically disordered regions (IDRs) tend to make promiscuous molecular interactions when their concentration is increased, and have been associated with dosage sensitivity [68]. We found that genes whose proteins have a high percentage of IDRs were more toxic for non-essential, but not essential, genes (**Figure S7A**, **Figure S7B, Table S6C**). Moreover, the toxic gene distribution was significantly skewed towards larger proteins for non-essential genes (KS test, p < 2 × 10^−14^, **Figure S7C**) but not for essential genes (KS test, p = 0.38, **Figure S7D, Table S6C**).

Finally, we compared our data with two genome-wide tests of overexpression toxicity that used either the barFLEX [11] or the GST-tagged [10] collection, both of which employ a GAL-inducible promoter for gene overexpression (**Figure S8A**). The GST-tagged collection contains strains expressing an NH_2_-GST tagged to the ORF in a high copy plasmid, while the barFLEX collection contains strains expressing untagged ORFs on a low-copy plasmid. To compare screens, we examined only those strains tested in all three studies, which included 608 Z_3_EV-toxic strains, providing a significant overlap with genes identified in the previous whole genome overexpression studies (χ^2^ test, p = 1.87 × 10^−19^, **Figure S8A, Table S6D**).

Overall, we observed a 50% overlap of toxic genes between the YETI collection and barFLEX collection and 30% overlap between YETI and GST-N strains. The larger overlap between barFLEX and YETI may reflect the fact that both collections involve untagged genes. In total, 346 out of 608 of Z_3_EV-toxic genes were only identified as toxic with the YETI collection (**Table S6D**). Finally, 27% of universally toxic genes (toxic in YETI, barFLEX, and GST) and 52% of Z_3_EV-specific toxic genes *only* showed growth impairment at the highest tested β-estradiol concentration (100 nM, **Figure S8B, Table S6E**). In other words, a smaller proportion of universally toxic genes *require* the highest level of inducer to show growth impairment as compared with Z_3_EV-specific toxic genes. Our ability to detect more growth defects with the Z_3_EV collection may be due to a combination of more precise expression control (a titratable promoter and an integrated test gene), as well as use of time-lapse quantification of growth.

### Application #1: Pooled BAR-seq screens

One of the design features of the YETI collection is the inclusion of unique barcodes marking each Z_3_EV promoter allele, enabling pooled growth assays, which use a barcode-based sequencing readout to assess competitive growth (BAR-seq) [69,70] (**Figure 5**). Competitive growth assays are extremely useful for quantifying growth phenotypes associated with mutation of non-essential genes. To test the YETI-NE collection for genes sensitive to under- and over-expression in liquid culture, we inoculated pools into YNB or SC medium with either 0 nM or 100 nM β-estradiol, under competitive growth conditions, for 48 hours in YNB or 36 hours in SC. Cultures were re-diluted at successive timepoints to maintain exponential growth and the SC timepoints were chosen such that the number of cell doublings would be similar to those achieved in YNB-grown cultures. At each timepoint, cells were first harvested and then the barcodes were PCR-amplified and quantified via next generation sequencing. The resulting data were normalized to data collected at time 0, then hierarchically clustered to look for general trends (**Table S7A**). This analysis revealed 11 clusters of strains which shared patterns of growth depletion dependent on the condition (YNB or SC) and/or the presence of β-estradiol (labeled in **Figure 5**). To identify potential biological explanations for the observed growth profiles, we performed functional enrichment analysis on the strain clusters. One cluster, Cluster YNB, was composed of strains depleted in 0 nM β-estradiol in YNB only. As described earlier, reduced growth in YNB and not SC is suggestive of auxotrophy. Indeed, we saw strong GO enrichment for biosynthetic processes, including amino acid and organic acid biosynthesis, and the average Aux score for genes in this cluster was relatively high (**Table S7B**).

**Figure 5:**
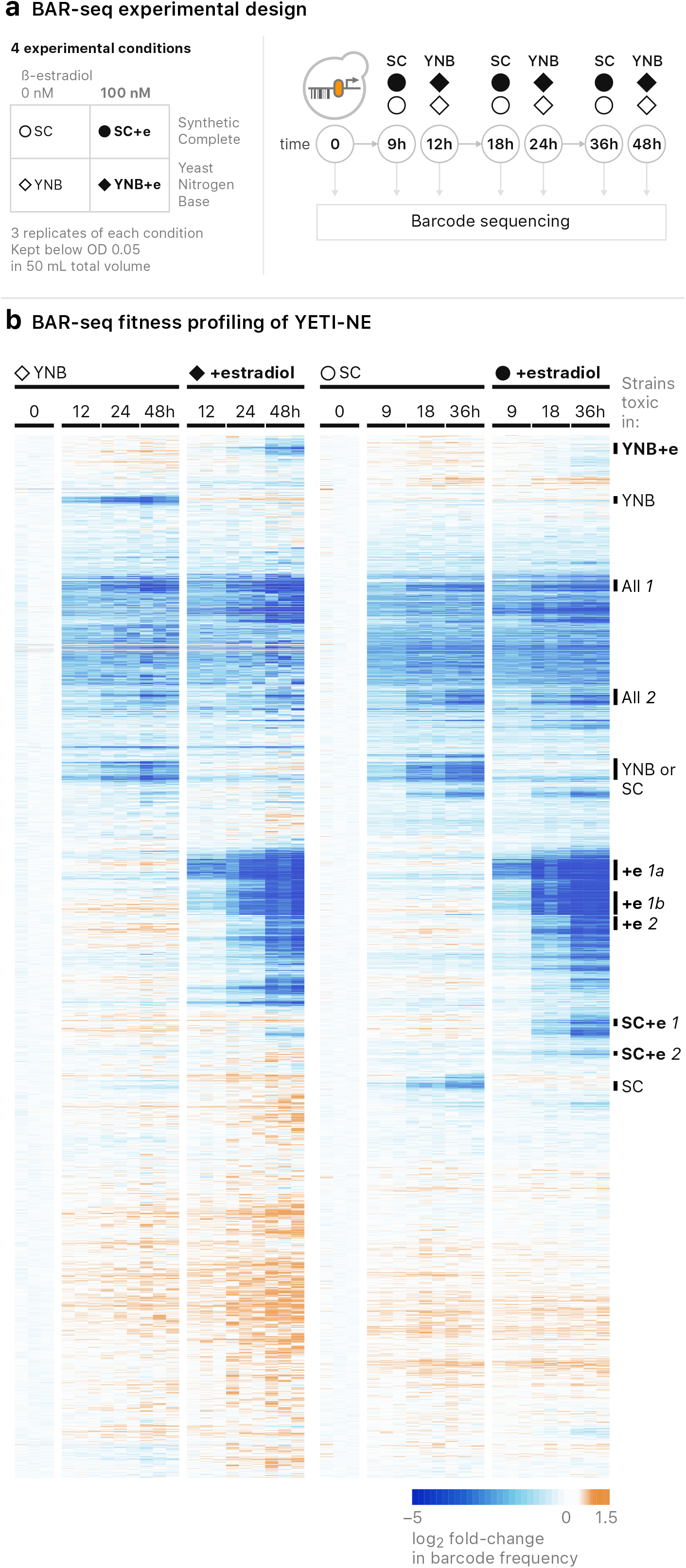
BAR-seq of YETI-NE strains. **A.** Diagram illustrating the pooled culture growth and harvesting strategy for BAR-seq experiments with YETI-NE strains. The labels in the diagram are also used to indicate the relevant growth conditions in the corresponding clustergram (part B). **B.** BAR-seq analysis of YETI-NE pools in SC and YNB medium in the absence or presence of 100 nM β-estradiol. Data were hierarchically clustered and 11 clusters labelled based on conditions in which strains exhibited reduced growth: Cluster YNB +e (showing depletion in 100 nM β-estradiol in YNB only), Cluster YNB (depletion in 0 nM β-estradiol in YNB only), Clusters All 1 and All 2 (time-dependent depletion in all conditions), Cluster YNB&SC (depletion in YNB and SC at 0 nM β-estradiol), Clusters “+e 1a” and “+e 1b” (fast depletion in YNB [12 h] and SC [9 h] at 100 nM β-estradiol), Cluster “+e 2” (depletion by 24 h in YNB and 18 h in SC at 100 nM β-estradiol), Clusters “SC+e 1” and “SC+e 2” (depletion in SC only at 100 nM β-estradiol), Cluster SC (depletion in SC only at 0 nM β-estradiol). Experiments were performed in biological triplicate.

Two other clusters were composed of strains that grew more poorly in the absence of β-estradiol, either only in SC medium (Cluster SC) or in both SC and YNB (Cluster “YNB or SC”). YETI-NE strains that became depleted over time in 0 nM β-estradiol, but not when their expression was induced, may encode proteins whose expression is important for growth. Consistent with this, both clusters were enriched for genes involved in cytosolic translation (13.5-fold enrichment for Cluster “YNB or SC”, p = 2×10^−34^ and 9.9-fold for Cluster SC, p = 5.9×10^−8^) and ribosome biogenesis (10.9-fold for Cluster “YNB or SC”, *p* = 1.8×10^−28^ and 8.3-fold for Cluster SC, p = 4.8×10^−7^), processes that are important for growth. The average single-mutant fitness was 0.65 for genes in Cluster YNB and SC and 0.72 for Cluster SC ([71], **Table S7B)**. We also found that genes in these clusters had higher native expression levels, on average, than all YETI-NE genes (**Table S7B**), consistent with their β-estradiol-dependent growth.

Changes in fitness in culture and on plates were positively and significantly correlated (Spearman correlation = .29 at 100 nM β-estradiol, p-value < 2.2 × 10^−16^) (**Figure S9**). Additionally, genes identified as toxic upon overexpression on plates showed the strongest agreement with clusters composed of strains that were depleted from the pooled culture in a β-estradiol concentration-dependent manner (Clusters “+e 1a”, “+e 1b”, and “+e 2”; 96%, 94%, and 72% overlap, respectively; **Table S7B**). Because we see quantitative agreement between data collected on plates and with BAR-seq, the ease of BAR-seq assays provides a robust paradigm by which the YETI collection can be used for screens in a highly parallel manner.

### Application #2: Mapping transcriptional and genetic interaction networks with YETI

Systematic analysis of transcriptome profiles in yeast strains whose growth is impaired by TF overexpression has been used to discover TF target genes [72]. To explore this approach with the YETI collection, we chose *ROF1*, which encodes a putative transcription factor containing a WOPR DNA-binding domain and whose induction with β-estradiol inhibits growth, both on plates and in BAR-seq experiments [73,74] (**Figure 5, Figure 6, Table S6A, Table S7A**). Overexpression of *ROF1* prevents “fluffy” colony morphology, a proxy for biofilm formation, in the F45 background; hence its name, Regulator of Fluffy [74]. Deletion of *MIT1*, the paralog of *ROF1*, results in a pseudohyphal growth phenotype in the Σ1278 genetic background, whereas a *rof1Δ* mutant has no phenotype in this assay [75].

**Figure 6:**
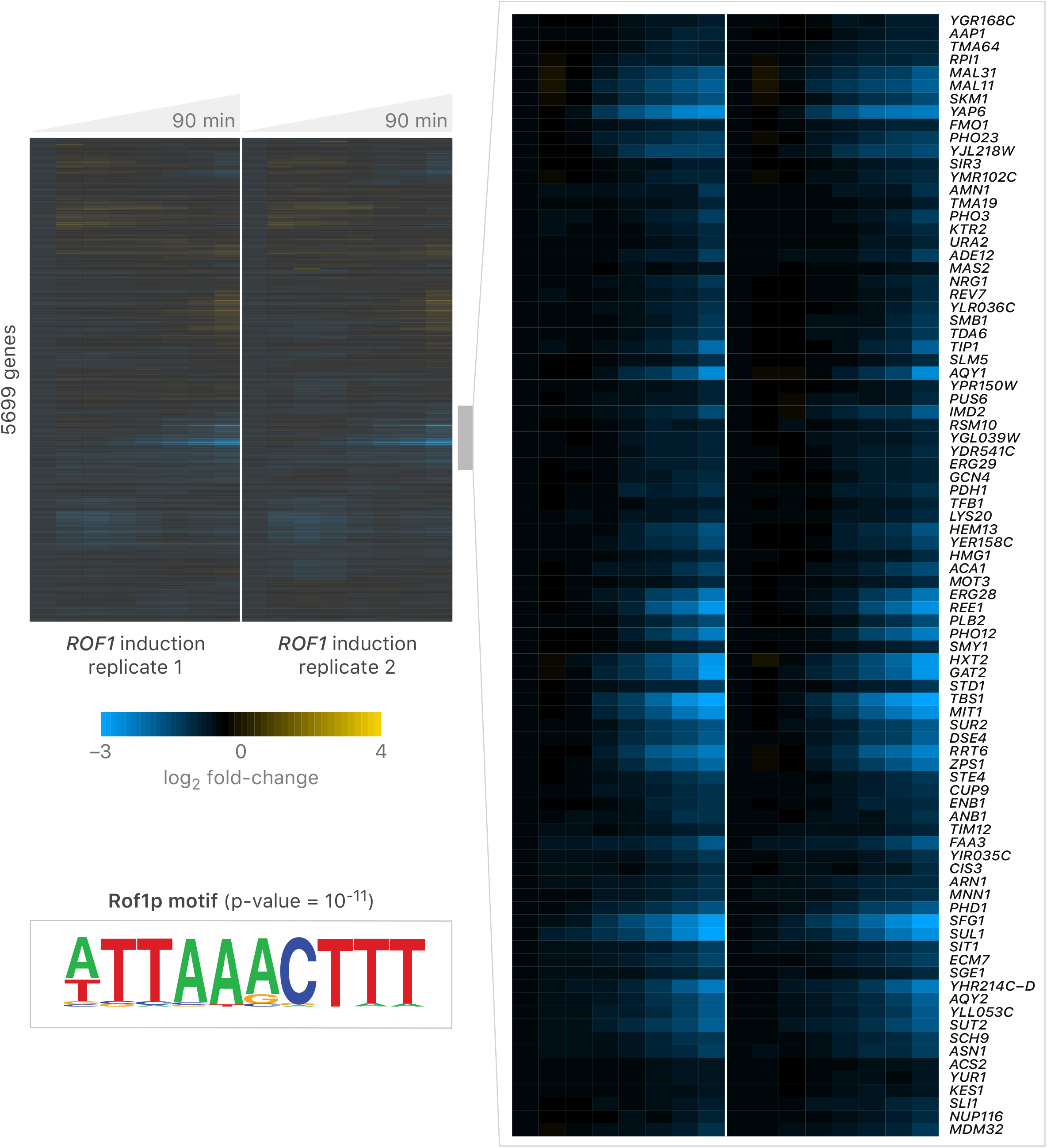
Rof1 is a transcriptional repressor. Microarray expression data resulting from induction of Z_3_pr-*ROF1* over a 90-minute time-course. The Z_3_pr-*ROF1* strain was grown in the presence of 1 μM β-estradiol in a phosphate-limited chemostat, and samples were harvested at the indicated times. One of two independent time courses is shown. Gene expression data were clustered using a Pearson correlation distance metric. One regulon of Rof1-repressed genes was identified, which is highlighted on the right. Motif enrichment was determined using the HOMER software suite [93].

The growth phenotype associated with *ROF1* overexpression in the Z_3_pr-*ROF1* strain provided an opportunity to explore the *ROF1* regulon and to illuminate *ROF1* function. We grew the Z_3_pr-*ROF1* strain to steady state in a phosphate-limited chemostat, induced *ROF1* with 1 μM β-estradiol, and tracked genome-wide gene expression changes over 90 minutes, as described previously [16,48]. The *ROF1-*YETI strain has less Rof1 than a WT strain does at t = 0 min. Upon induction with β-estradiol, in duplicate experiments, we observed a subset of genes whose expression was rapidly reduced upon *ROF1* overexpression, suggesting Rof1 acts as a transcriptional repressor (**Figure 6, Table S8A**). Using HOMER (Hypergeometric Optimization of *Motif* EnRichment; [76]), we found that the promoters of genes repressed by Rof1 are most strongly enriched for a WOPR-domain-like DNA-binding sequence, [A/T]TTAAACTTT (p-value ~ 10^−11^). The Rof1-repressed genes were enriched for several functional processes, including water transport, siderophore transmembrane transport, and repressing transcription factor binding activity (**Table S8B**). Several TF genes were repressed by Rof1, including *MOT3*, *CUP9*, *PHD1*, *YAP6*, *NRG1*, *GAT2*, *TBS1*, *ACA1*, *GCN4, CIN5*, and a gene encoding Rof1-paralog, *MIT1*. Our initial analysis of *ROF1* indicates that it encodes a *bona fide* TF that is highly interconnected with other TFs and illustrates the usefulness of ZEV-promoter strains for exploring TF function.

Synthetic dosage lethal (SDL) interactions, which occur when overexpression of a gene causes a more severe fitness defect when a second gene is mutated, can reveal information about pathways and bioprocesses affected by the overexpression. YETI strains carry SGA markers, enabling introduction of Z_3_EV induction alleles into appropriately marked arrays of yeast mutant strains. We used this feature to further explore the genetic basis of the fitness defect caused by *ROF1* overexpression. We crossed a query strain containing Z_3_pr-*ROF1* with the deletion collection and the collection of temperature-sensitive alleles of essential genes in the absence of β-estradiol, and then we induced *ROF1* overexpression with 1 μM β-estradiol at the final SGA selection step. We discovered 264 unique genes that had SDL interactions with *ROF1* (**Table S8C**), with the strongest interactions involving members of the cell wall integrity (CWI) pathway, *SLT2*, *BCK1* and *PKC1*, and two genes with roles in cell wall biosynthesis, *GFA1* and *QRI1*, consistent with a role for *ROF1* in surviving cell wall stress [77]. *ROF1* also showed SDL interactions with numerous genes involved in chromatin remodeling and general transcription, including multiple members of the Swr1 complex, Ino80 complex, Rpd3L complex, NuA4 complex, and COMPASS, which presumably reflects its role as a transcriptional repressor and a regulator of other TFs. We conclude that overexpressing *ROF1* causes a stress response making cells dependent both on the CWI pathway and on multiple transcription pathways for normal growth.

### Reversibility of the Z_3_EV promoter alleles

Our experiments with a Z_3_EV-GFP allele showed that the Z_3_EV promoter could be both induced by the addition of β-estradiol and repressed upon its removal (**Figure 1C**). This is an important feature that could prove particularly useful for the study of essential genes. We examined whether expression of the Z_3_EV promoter was more generally reversible by assaying the growth of strains carrying Z_3_EV promoter alleles of essential genes. We chose 24 Z_3_EV strains whose growth was β-estradiol-dependent, sporulated the Z_3_EV diploids, and then selected haploids containing the Z_3_EV system in the presence of 10 nM β-estradiol, which allowed good growth for all the strains. We then transferred the strains to the same medium containing 0 nM β-estradiol or 10 nM β-estradiol, incubated for two days, and then repeated the transfer and growth (**Figure S10**, **Table S9**). Three of 24 strains failed to grow on 0 nM β-estradiol, showing strong reversibility of the Z_3_EV promoter allele, and another 5 strains had Z_3_EV promoter alleles that showed partial reversibility in this assay. However, 16 of the 24 (67%) Z_3_EV strains grew in the absence of β-estradiol, even though their initial growth had been β-estradiol-dependent. These data suggest that many essential genes under the control of Z_3_EV may continue to be expressed in the absence of β-estradiol for numerous cell doublings.

We also examined reversibility of overexpression toxicity (see Methods for details). We chose 19 strains from the YETI-NE panel carrying Z_3_EV alleles that caused at least a ~50% growth defect in the presence of 100 nM β-estradiol (**Figure S11A**). These strains were first pinned onto SC lacking β-estradiol, and then colonies were transferred to fresh YNB plates either with or without β-estradiol. After ~24 hr of growth, colonies were transferred again to YNB plates with or without β-estradiol to initiate a 120-hr time lapse growth assay. Of the 19 toxic Z_3_EV promoter alleles tested, 13 (68%) were fully or partially reversible. For example, the Z_3_EV*-TIP41* allele showed a strong reversibility phenotype (**Figure S11B**); when pinned from 10 nM to 0 nM β-estradiol, growth is largely restored to normal levels (**Figure S11C**). In contrast, the Z_3_EV*-SGS1* allele caused toxicity that was not reversible (**Figure S11D**). Although we have only examined a subset of Z_3_EV promoter alleles for reversibility, our analyses suggest that overexpression toxicity phenotypes associated with Z_3_EV promoter allele expression are more easily reversed than growth phenotypes caused by removing β-estradiol from the growth medium.

### Z_3_EB42: a reversible artificial transcription system for lowly expressed genes

Our characterization of the Z_3_EV system revealed limitations, including the growth of many strains expressing Z_3_EV promoter alleles of genes with low endogenous expression even in the absence of inducer (**Figure 2**). Previous work has shown that changing the number of Zif268 binding sites in the β-estradiol-dependent promoter, and/or modifying the activation domain of an artificial transcription factor (ATF), can expand the range of possible gene expression levels [14,78]. Therefore, we re-engineered Z_3_EV as well as its target promoter, Z_3_pr, with the goal of designing a system with reduced ‘leakiness’ of gene expression in the absence of inducer. Specifically, we engineered promoters with different numbers of Zif268 binding sites (6 or 2) (**Figure 7**, **Figure S12**). We also employed the use of yeast regulatory elements to attempt to further reduce the strength of Z_3_pr [40]. Between position −158 and −146 of the *CAR1* gene is an upstream repression sequence (URS1), required for *CAR1* repression [79]. Ume6, a DNA binding protein, binds to the URS1 sequence and recruits the Sin3-Rpd3 complex for transcriptional repression [80]. Ume6 represses sporulation-specific genes during mitotic growth and genes involved in arginine catabolism when readily catabolized nitrogen sources are available in the environment [82] so we anticipated that it would have repression activity under our standard conditions. Thus, we appended a region extending from nucleotides −198 to −1 of *CAR1* to the 3’-end of Z_3_pr (6x or 2x Zif268 binding sequences [bs]). Finally, we created a new ATF, Z_3_EB42, with a B42 transcriptional activation domain in lieu of VP16. B42, a short unstructured acidic peptide encoded by an *Escherichia coli* genomic DNA fragment, has weaker activity than VP16 [78,83].

**Figure 7:**
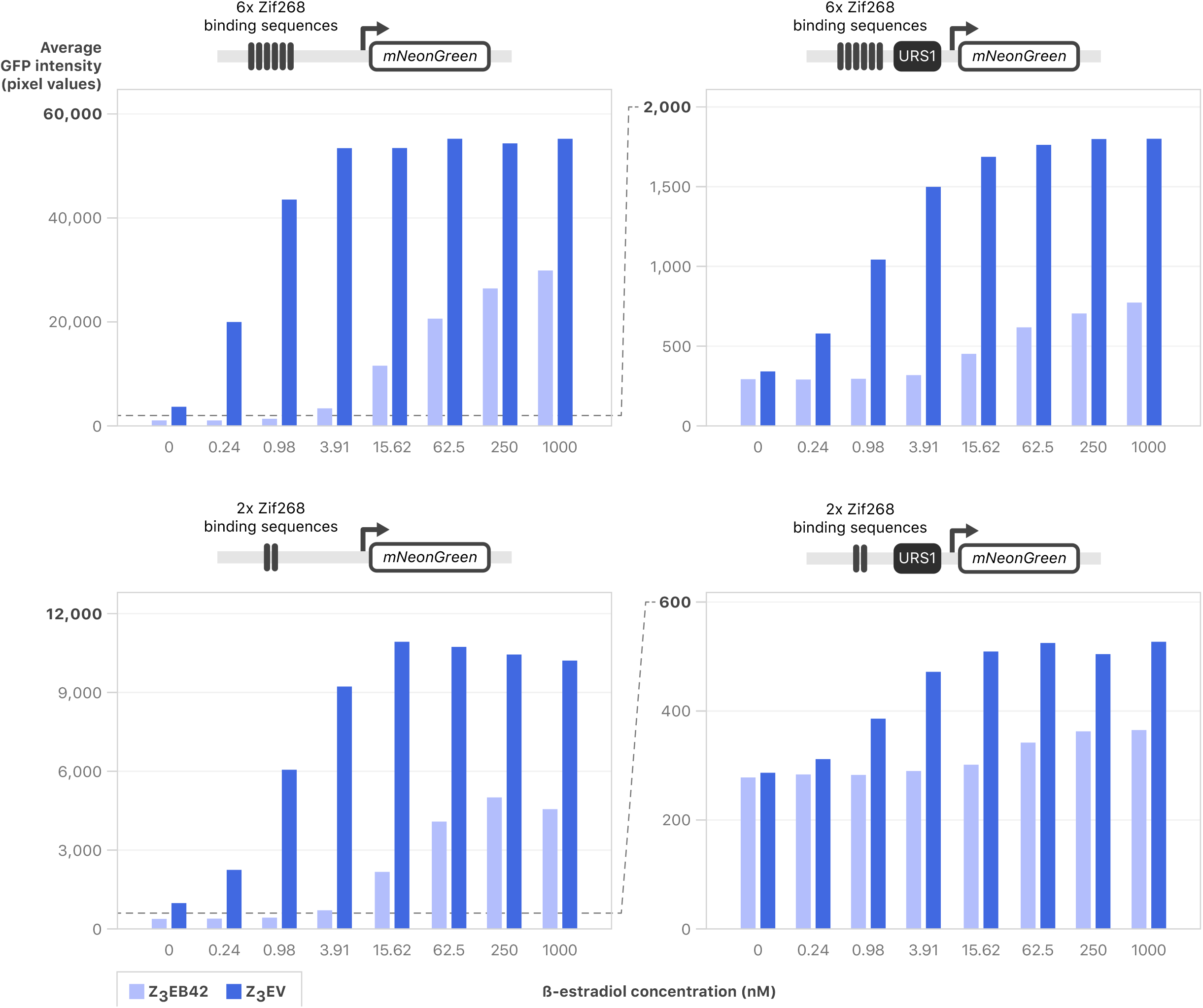
Number of binding sites, URS1 presence, and ATF choice all affect mNeonGreen reporter induction. Strains with all combinations of ATFs and promoters driving mNeonGreen were grown to early log phase in YNB and mNeonGreen was induced with various levels of β-estradiol. Strains were imaged every hour using a Phenix automated confocal microscope and average fluorescence/cell was calculated using Harmony software. Average of two replicates is shown at t = 8 hours following induction with >300 cells quantified per dose. Full time series are shown in **Figure S12**.

We constructed eight combinations of ATFs and Z_3_ promoters and quantified dose-response curves across a range of β-estradiol concentrations using an mNeonGreen reporter (**Figure 7**, **Figure S12**). As predicted, URS1 had a strong effect on mNeonGreen production, resulting in a 20-fold decrease in production for Z_3_EV and a 14-fold decrease in production for Z_3_EB42 for 2x Zif268 bs in high β-estradiol. (**Figure 7**). Additionally, at saturating levels of inducer, we saw that the use of Z_3_EV resulted in ~2-fold more fluorescent signal than Z_3_EB42 for each promoter. Consistent with previous work, reducing the number of Zif268 binding sites in Z_3_pr reduced fluorescent signal [14] (**Figure 7**). The weakest promoter in our panel was Z_3_(2x Zif268 bs + URS1)pr, which we refer to as Z_3_pr-Zif2-URS1. The combination of Z_3_EB42 and Z_3_pr-Zif2-URS1 resulted in the least leaky expression in the absence of inducer, as well as the lowest mNeonGreen signal. Finally, we assessed the reversibility of each ATF-promoter pair, and we found that URS1 made an important contribution to reversibility: for both Z_3_EV and Z_3_EB42, only promoters containing URS1 were able to reverse expression to background levels in <24 hours (**Figure S13**). Z_3_EB42 and Z_3_pr-Zif2-URS1 were the most reversible ATF-promoter pair. We now refer to the combination of the ATF, Z_3_EB42, and the synthetic promoter, Z_3_pr-Zif2-URS1, as the Z_3_EB42 system (**Figure 8A**).

**Figure 8:**
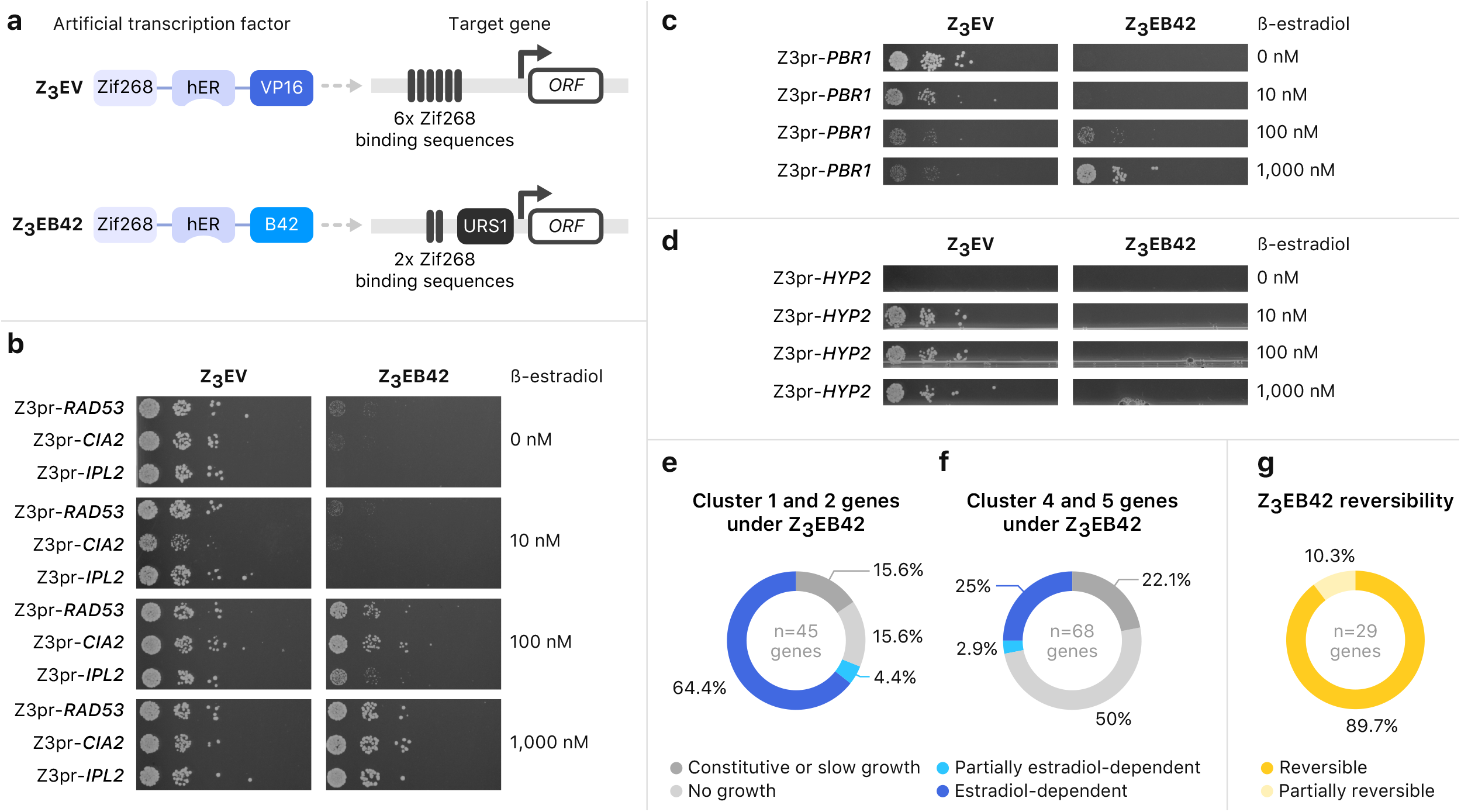
Engineered gene expression system for more stringent regulation. **A**. Diagrams of Z_3_EV artificial transcription factor with Z_3_pr promoter and Z_3_EB42 artificial transcription factor with Z_3_pr-Zif2-URS1 promoter. **B-D.** Serial spot dilutions onto medium containing various concentrations of β-estradiol. Diploid strains containing the indicated regulatory systems (see A) controlling the target genes were incubated in sporulation medium for 5 days. Serial dilutions of cells were plated on SD-his-ura-arg-lys with canavanine, thialysine and ClonNAT with indicated concentrations of β-estradiol to obtain haploid cell growth. **E.** Distribution of growth phenotypes for strains carrying Z_3_EB42 promoter alleles for 45 essential genes from Cluster 1 and 2 (see Figure 2). The color key for interpreting the figure is below panels e-g. **F.** Distribution of growth phenotypes for strains carrying Z_3_EB42 promoter alleles for 68 essential genes from Cluster 5 (see Figure 2). **G.** Distribution of Z_3_EB42 promoter reversibility phenotype for 29 tested essential genes.

We compared Z_3_EV and Z_3_EB42 function on promoter replacement alleles of a panel of essential genes: three that enabled growth without β-estradiol (*RAD53*, *CIA2*, and *IPL2*), one that allowed growth without inducer but was toxic when overexpressed (*PBR1*), and one that gave β-estradiol-dependent growth (*HYP2*) with the Z_3_EV system. Strains with the Z_3_EV system driving *RAD53*, *CIA2* or *IPL2* grew in the absence of β-estradiol (**Figure 8B left**). Strains with these genes under the control of Z_3_EB42 were not able to grow in 0 or 10 nM β-estradiol, indicating that any leaky expression was repressed with the stringent system (**Figure 8B right**). *PBR1* was leaky and had a growth defect on 1000 nM β-estradiol when regulated by the Z_3_EV system (**Figure 8C left**). In contrast, a strain with Z_3_EB42 system driving *PBR1* expression did not grow in the absence of β-estradiol but grew in 1000 nM β-estradiol (**Figure 8C right**). The growth defect phenotype caused by *PBR1* overexpression in 1000 nM β-estradiol with the Z_3_EV system was not detected with Z_3_EB42. Finally, we tested a Z_3_EB42-inducible allele that behaved well in the YETI allele collection, with no growth in the absence of β-estradiol and normal growth in the presence of 10 nM or higher β-estradiol (**Figure 8D left**). The Z_3_EB42-regulated *HYP2* did not support growth even at 1000 nM β-estradiol (**Figure 8D right**). These results indicate that, for some essential genes, Z_3_EB42 leads to levels of gene expression low enough that growth is β-estradiol-dependent, but for others, it may not enable enough expression to support growth and may not cause phenotypes that depend on overexpression.

Since we found that the Z_3_EB42 system conferred β-estradiol-dependence for some essential genes that were not regulated with Z_3_EV, we expanded our survey of Z_3_EB42 system performance to include a larger set of essential genes that had given either β-estradiol-dependent (Clusters 4 and 5) or β-estradiol-independent (Clusters 1 and 2) growth with the Z_3_EV system. Of 45 strains with genes that were β-estradiol-independent, 7 (15.6%) gave constitutive growth, 29 (64.4%) were β-estradiol-dependent, 2 (4.4%) were partially β-estradiol-dependent and 7 (15.6%) gave no growth when controlled by Z_3_EB42 instead of Z_3_EV (**Figure 8E, Table S10**). Of 68 strains with genes that were β-estradiol-dependent with Z_3_EV, 15 (22%) were constitutive, 17 (25%) were β-estradiol-dependent, 2 (3%) were partially β-estradiol-dependent and 34 (50%) were not able to grow at any β-estradiol concentration when controlled by Z_3_EB42 (**Figure 8F, Table S10**). Because the genes from Clusters 4 and 5 tend to have higher native transcript levels, it is likely that the Z_3_EB42 system does not drive sufficient gene expression to support growth for a significant fraction of highly expressed essential genes. Finally, we asked if the lower-expression Z_3_EB42 system could confer reversible growth. Of 29 genes from Cluster 1 and Cluster 2 that had β-estradiol-dependent growth with Z_3_EB42, we found 26 (90%) were reversible and 3 (10%) were partially reversible (**Figure 8G**). These data demonstrate that the Z_3_EB42 system confers high reversibility of essentiality and provides a useful tool for investigating the loss-of-function of Cluster 1 and 2 genes.

We conclude that the Z_3_EV system can achieve conditional growth for many essential genes, and that the Z_3_EB42 system tends to enable conditional (and reversible) growth in cases where Z_3_EV cannot. Thus, when growth is the readout, the best choice of expression system depends on the individual properties of a specific gene.

## Discussion

We constructed and characterized YETI, a genome-scale collection of Z_3_EV inducible alleles of yeast genes. Previously, we established that combining inducible expression with transcriptome-wide time series measurements is an effective strategy for elucidating gene regulatory networks. The ability to “switch” a gene on and measure dynamically how every other gene responds has allowed us to identify causal regulatory interactions, both direct and indirect, and observe instances of feedback control that were previously inaccessible [16,48]. By creating the YETI collection, we are expanding the uses of this system from studies of individual genes to nearly all genes in the yeast genome. To characterize the YETI collection and provide information to guide its use by the community, we have performed quantitative growth-based phenotyping, and carried out pooled BAR-seq screens. To demonstrate the utility of this YETI collection, we mapped the regulon of a putative TF and identified its SDL interactions. Collectively, these experiments validated the quality of the library, revealed new biology, illuminated an important though unappreciated design principle of synthetic gene control, and motivated the development of the Z_3_EB42 system for applications requiring more refined regulation.

Because our YETI strains were constructed first as diploids, we harnessed the power of yeast genetics to sporulate and select haploid strains with one copy of Z_3_pr-controlled essential genes at differing levels of inducer. This unique experimental design revealed that ~40% of Z_3_pr-controlled essential genes had enough transcript to facilitate growth in the absence of inducer, even though Z_3_pr is extremely weak. Many genes in bacteria, yeast, and humans are lowly expressed with only a handful of transcripts per cell [84]. Making a strong inducible system that is also completely “off” in the absence of β-estradiol is an engineering challenge. Indeed, the cost of achieving strong inducible expression is often some basal level of leakiness, which can be problematic for growth-based screens. To determine the extent of this problem, in yeast and *E. coli*, we took advantage of a ground-truth set of essential genes.

Unlike yeast and *E. coli*, we have yet to define a clear-cut ground-truth dataset of essential genes in human cells. Many human cell essential genes have been identified in a variety of different cell lines from pooled-screens, and we find that annotated essential genes are ~10x-20x more highly expressed than those defined as non-essential (**Figure S14**). In yeast, essential genes are, on average, expressed only ~1.4x more highly than non-essential genes. The difference in expression between essential and non-essential genes could be much greater in human cells than it is in yeast, or pooled screens in cell lines may be sampling the most extreme parts of the expression distribution. We favor the latter interpretation and believe that the loss of highly-expressed essential genes is simply easier to detect than the loss of lowly expressed essential genes with currently available technology. This interpretation also implies that CRISPR gene inactivation approaches are more effective at disrupting highly expressed genes than lowly expressed ones. Mechanistically, low levels of chromatin accessibility, which is associated with low levels of gene expression, could prevent both Cas9 and dCas9 from accessing specific gene targets. Focusing on yeast and *E. coli*, we found that generating low expression of already lowly-expressed essential genes, using multiple approaches, including Z_3_EV, Tet-off, DAmP, TS-degron and CRISPRi, is problematic for growth-based screens. In yeast, recently improved Tet-based systems with low expression variation and reduced leakiness may be useful for some applications, including expressing lowly expressed genes, as well as multiplexed experiments with the YETI collection [85,86]. Here, we developed Z_3_EB42 to specifically enable conditional and reversible growth with lowly expressed essential genes.

The YETI collection provides new opportunities to elucidate regulatory interactions and instances of feedback control on a genome scale. We anticipate that the barcode feature will enable this collection to be used for highly-parallel pooled screens [69]. Indeed, our collection provides an opportunity to perform pooled screens without plasmids and at many different levels of expression. Furthermore, the YETI collection provides a new and general system for functional analysis. Of particular interest to us is that replicative aging in yeast is a model for studying the aging process in eukaryotes. For example, it should be possible to combine the YETI collection with Miniature-chemostat Aging Devices (MADs) [87] to explore how molecular networks change with age. MADs give us the ability to obtain replicatively aged cells that are rare in standard cultures. Combined with YETI, we anticipate that we can explore how titrating individual genes affects the aging process itself. We also believe that there are opportunities in metabolic engineering and industrial biotechnology applications for titrating target genes to increase the yields of useful products. To make the YETI collection compatible with perturb-Seq-like approaches for monitoring regulatory networks at the single-cell level, barcodes encoded in the RNA could be introduced [88]. Finally, each strain in this collection contains the SGA markers, which means future strain engineering can be done using automated approaches. The combination of YETI with fluorescent markers will enable phenotypic screens that can be performed in pooled or arrayed formats.

## Materials & Methods

### Strain and plasmid construction

The yeast strains and plasmids used in this study are listed in the **Supplement**. To construct the parental diploid yeast strain Y14789 expressing the Z3 binding domain-ER-VP16 transcription factor (Z_3_EV) and carrying SGA markers, a DNA fragment containing a *natMX*-marked Z_3_EV fragment was PCR amplified from strain DBY12394 using primers that contained sequences that enabled integration into genome downstream of the repaired *HAP1* locus in the strain RCY1972 (gift from Dr. Amy Caudy). The strain was then crossed to Y7092 to create a diploid strain, and Y14851 was isolated by tetrad dissection. Subsequently, Y14851 was crossed to Y14537 to generate diploid strain Y14789, which was used for construction of the β-estradiol-inducible allele strain collection. To construct the source plasmid for Z_3_EVpr (p5820), the kanMX marker in plasmid pRB3564 [13] was replaced with the *S. cerevisiae URA3* gene. The Z_3_ promoter (Z_3_pr) is a derivative of the *GAL1* promoter in which a region containing three canonical *GAL4* binding sites (5’-CGG-N_11_-CCG-3’) is replaced with six Zif268 binding sites (5’-GCGTGGGCG-3’). In creating p5820, we also removed a non-canonical Gal4 binding site from pRB3564. The sequence of the Z_3_EV transcription factor coding region was previously described. The plasmids p7418 and p7419 were constructed by modifying p5820.

To construct β-estradiol promoter replacement alleles for each *S. cerevisiae* gene, the Z_3_ synthetic promoter linked to *URA3* was PCR-amplified from plasmid p5820. The forward primer contained homology to 40 bp upstream of the ATG of each gene with a unique 12 nucleotide barcode. The reverse primer contained 40 bp complementary to the region immediately downstream of the ATG (**Table S2A, B, C**). PCR products were transformed into strain Y14789 and transformants were selected on SC-ura+ClonNAT at 30°C. The integration locus was confirmed by colony PCR, as previously described [27]. Candidate clones were streaked onto YPD+ClonNAT plates, and single colonies were isolated and replica-plated onto SC-ura to check the stability of the integrated marker. For non-essential ORFs, we also constructed a *MAT***a** haploid collection, by sporulating the relevant diploid strains for 5-6 days. Following sporulation, single colonies were isolated by streaking the spore mixture onto SGA selection medium SC-his-ura-arg-lys with canavanine, thialysine and ClonNAT [28] and the plates were incubated at 30°C for 4 days. The integration locus was re-confirmed by colony PCR. The mating types and selectable markers of the haploid strains were also confirmed. To make Y15292, a GFP fragment and the Z_3_ synthetic promoter were PCR amplified then integrated into the *ho* locus of Y14851. For testing the stringent system, Z_3_EB42 was integrated downstream of the repaired *HAP1* locus. To make plasmid p7418, the six Zif268 binding sites of Z_3_EVpr (p5820) were replaced with two Zif268 binding sites. One hundred and ninety-seven base pairs of *CAR1* upstream sequence including URS1 was appended to the 3’ end of the promoter (**Supplement**).

The YETI-E and YETI-NE diploid collections have the genotype: *MAT***a**/α *ORF*+/[*barcode::URA3::Z3EVpr-ORF*] [*HAP1::natMX-ACT1pr-Z3EV-ENO2term*]/*HAP1 ura3Δ0*/*ura3Δ0* [*can1Δ::STE2pr-Sphis5*]/*CAN1 his3Δ1*/*his3Δ1 lyp1Δ*/*LYP1*. Following selections, the YETI-E and YETI-NE haploid genotypes are *MAT***a**[*barcode::URA3::Z3EVpr-ORF*] [*HAP1::natMX::ACT1pr-Z3EV-ENO2term] ura3Δ0 can1Δ::STE2pr-Sphis5 his3Δ1 lyp1Δ*

### Assessing expression of GFP kinetics using flow cytometry

To assess the dependence of gene expression on the concentration of β-estradiol in the experiments shown in Figures 1 and S1, the yeast strain Y15292, carrying a GFP reporter gene driven by Z_3_EVpr, was incubated overnight at 30°C in minimal medium. The overnight culture was then diluted into fresh medium and grown at 30 °C to a density of 6 × 10^6^ cells/mL in the presence of various concentrations of β-estradiol. To measure the kinetics of expression, cells were grown as described above but with medium containing 10 nM β-estradiol (Fig 1D). To measure the effect of removing β-estradiol on GFP expression, cells were washed in PBS 3 times, then incubated in fresh medium without β-estradiol. At each timepoint, cells were harvested and fixed in 70% ethanol. Subsequently, cells were washed with 1 ml PBS 3 times and 5 μg/ml propidium iodide was added. To measure GFP signal in single yeast cells, we used fluorescence-activated cell sorting to record 10,000 events using a BD FACS Aria lllu with a high throughput sampler (BD Biosciences). We gated for single cells, which were identified by analyzing the signal width of the forward and the side scatter. Dead cells, which were stained with propidium iodide, were excluded from data analysis.

### Z_3_(2 binding sites) URS1 promoter alleles of essential genes

Growth of essential gene alleles was assessed by sporulating the heterozygous diploid strains on sporulation plates for 5 days and then either spotting (with 10x dilutions) or directly pinning onto SC-his-ura-arg-lys with canavanine, thialysine and ClonNAT plates in the absence or presence of various concentrations of β-estradiol.

### Preparation of Agar Plates for Growth Profiling of YETI-E and YETI-NE strains

All the agar plates were prepared by either Universal Plate Pourer (KREO Technologies Inc.) or manually in Nunc™ OmniTray™ Single-Well Plate (ThermoFisher Scientific 264728). To minimize noise of colony size, media volume in each plate was set to 40mL, and plates with air bubbles, particles, uneven surfaces, and scratches on the bottom of the plates were discarded. After agar was solidified, plates were flipped (upside down) and dried for 48 hours at room temperature to maintain the same moisture level in each plate.

### Pinning and Imaging of Agar Plates

Pinning of yeast strains was done with a BM3-BC pinning robot (S&P Robotics, Inc.) using various pin tools indicated below. Incubation of plates at 30℃ and imaging of plates were done using the spImage-A3 instrument (S&P Robotics, Inc.) with manual focus mode of camera setting. To reduce the time lag from the pinning and imaging of plates, less than 32 plates were processed at once, and imaging was started immediately after pinning on final assay plates with β-estradiol. Plate images were generated every hour up to the time point indicated in each experiment (mostly up to 60 hours).

### Image processing, quantification of yeast colonies, and growth phenotypes

For most of the reported data, colony sizes were quantified using Platometer, an open-source image-processing Python tool available at https://github.com/baryshnikova-lab/platometer. Briefly, Platometer automatically detects the grid of colonies on an image and estimates the size of each colony as the number of pixels above background after adaptive thresholding. Following quantification, raw colony sizes were normalized for plate effects, positional effects and competition effects as described previously [66]. Images were taken every hour for 60 hours, and normalized colony sizes across consecutive timepoints were assembled into growth curves. For each colony, the area under the growth curve (AUGC), which integrates the main three aspects of yeast population dynamics (duration of lag phase, exponential growth rate and carrying capacity), was used as a global estimate of growth efficiency.

### Growth Profiling of YETI-E Strains

YETI-E strains (1,022 heterozygous diploids) were re-arrayed to randomize positions of each strain on agar plates in 384 format. One row and one column on the edge was filled with wild type strain (Y15090) to exclude nutritional and spatial advantage for the strains on the edge. Two sets of the rearrayed (position-randomized) YETI-E collection were pinned again to generate quadruplicate 1,536 format, and multiple copies were prepared using a 1,536 0.8 mm pin tool. All the pinnings for re-arraying and copying were done on ‘Z_3_EV diploid maintenance plates’. The colonies were pinned onto ‘enriched sporulation plates’ for Z_3_EV diploid’ and incubated at room temperature for 5 days. Thereafter, sporulated colonies were pinned onto ‘Z_3_EV essential haploid selection with β-estradiol plates’ (see Supplement) with a 1,536 0.5 mm pin tool. 0.5 mm pins are the lower limit in size to deliver small amounts of yeast cells to destination plates to generate a larger dynamic range of growth, and not to puncture agar plates. Two technical replicates were processed for two position-randomized libraries at 12 β-estradiol concentrations (0, 0.01, 0.03, 0.1, 0.3, 1, 3, 10, 30, 100, 300, and 1,000 nM).

### Growth Profiling of YETI-NE strains

The non-essential haploid collection containing ~4,600 strains was processed in the same manner as the essential collection except that the wild type strain on the edge was different (Z_3_pr-*ho*), and the library came from the ‘Z3EV haploid maintenance plate’ (complete media), and was pinned onto SC or minimal media. Two sets of experiments were performed: pinning from complete medium to complete medium with β-estradiol and pinning from minimal medium to minimal medium with β-estradiol. Doses of β-estradiol were 0 nM, 1 nM, 5 nM, 10 nM, and 100 nM.

### Estimate of fraction of defective strains

Correct insertion of the URA3::Z_3_pr cassette was confirmed by PCR for all strains. To estimate the fraction of defective strains, i.e. strains where the Z_3_pr was not inducing expression of the gene, we used data from the YETI-E collection, for which growth as a haploid strain was a reflection of correct Z_3_pr insertion. Following our initial construction, we identified 90 strains that had little or no growth at all concentrations of β-estradiol. We were able to reconstruct 47/54 of these to give strains that had growth at some concentration of β-estradiol. From this we estimated that 47 strains were initially defective out of 986 (932 strains in Cluster 1, 2, 4 and 5, and 54 strains from our reconstruction), yielding a percentage of defective strains of 4.8%.

### Identifying genes that impair growth

Toxic strains were identified from the YETI-NE and YETI-E collections by growth on synthetic complete media at 0 (x) and 100 nM (y) β-estradiol concentration. We first normalized our y values for unexpected changes in growth due to the addition of β-estradiol. Strains at growth extremes, the top 2% and bottom 4% at 0 nM and 100 nM were removed. We expect the remaining × (0 nM) and y (100 nM) values to have a 1:1 relationship. We fit a linear model: y_measured_ = m*x_measured_+b. *y*_corrected_ = *x*_predicted_ (*x*_predicted_=(*x*_measured_-b)/m). We can then calculate changes in growth on β-estradiol using the distance from the line equation = abs(a*x*_measured_ +b*y*_corrected_ + c) / sq_root(a^2^ + b^2^). Genes with a distance from the diagonal larger than 2,000 were considered toxic and used in downstream analysis. This distance cut off agreed with genes identified as toxic from YETI-E clustering analysis.

### Reversibility of overexpression phenotypes

Strains were first pinned onto SC lacking β-estradiol, and then colonies were transferred to fresh YNB plates either with or without β-estradiol. After ~24 hr of growth, colonies were transferred again to minimal medium-containing plates with or without β-estradiol to begin a 120-hr time lapse growth assay. Four conditions were tested: 1) transfer from no β-estradiol to no β-estradiol (0-0), 2) transfer from 10 nM β-estradiol to 10 nM β-estradiol (continuous β-estradiol, 10-10), 3) transfer from no β-estradiol to 10 nM β-estradiol (addition of β-estradiol) and 4) transfer from 10 nM β-estradiol to no β-estradiol (removal of β-estradiol). Growth-curve AUCs were calculated for each strain in each condition.

### RNA-seq

To compare the expression of Z_3_pr-driven alleles to that of native promoter-driven alleles, we sequenced RNA from a subset of strains after induction with nine different doses of β-estradiol.

Strains were grown overnight in SC + ClonNAT with glutamate as a nitrogen source in a 96 deep-well plate. In the morning, cultures were diluted to an OD of 0.1 in the same media and the culture was grown for 5 hours until the cells reached log phase. Once the cells reached exponential phase, 1 μL of cells was taken and flash frozen in liquid nitrogen as an initial timepoint. β-estradiol was then added to the remaining cultures and the cells were grown for an additional 30 minutes before the final time points were taken. All samples were frozen in 7 μL of mastermix, which was comprised of 0.8 μL of 10x lysis buffer, 3.15 μL of dH_2_O, 1 μL of ERCC RNA spike ins, 0.05 μL of RNAse inhibitor, and 2 μL of oligo-dT primer (sequence: AAGCAGTGGTATCAACGCAGAGTACTTTTTTTTTTTTTTTTTTTTTTTTTTTTTTVN). The samples were then processed as follows:

1. <u>Reverse Transcription</u>: The samples were freeze-thawed 3x and incubated at 72°C for 3 minutes and 4°C for 2 minutes to denature the RNA secondary structure and enhance oligo-dT primer binding. 12 μL of the reverse-transcription mastermix (containing 4 μL 5x FS buffer, 1 μL 48 μM LNA template switch oligo (sequence: AAGCAGTGGTATCAACGCAGAGTACrGrG+G), 2uL 10mM dNTP mix, 2.5uL 20mM DTT, 0.5uL RNAse inhibitor, and 2 μL reverse transcriptase) was added to each sample and they were incubated at 42°C for 90minutes, 70°C for 10minutes, and kept at 4°C until the next step.
2. <u>cDNA amplification</u>: 30 μL of a PCR mastermix (25 μL 2x SeqAmp PCR buffer, 1 μL 12uM ISO PCR primer (sequence: AAGCAGTGGTATCAACGCAGAGT), and 1 μL seqAmp DNA polymerase, and 3 μL dH_2_O) was added to the sample. 10 PCR cycles were then run on the samples. 95C for 1 minute, 10x of 98°C for 10s, 65°C for 30s, 68°C for 3 min, 72°C for 10min, and then kept at 4°C until the next step.
3. <u>cDNA clean up</u>: an XP bead cleanup was done at 0.9x and 11 samples were run on the bioA to check quality and concentration. The rest of the samples were quantified using pico-green.
4. <u>Library prep and sample pooling</u>: The Nextera XT kit was used for library prep. 2 μL of each sample were pooled together and a double XP bead cleanup was done on the pooled libraries before running on the sequencer (hiseq 4000).
5. After demultiplexing, fastq files were processed using Salmon and aligned using STAR to obtain tpms.

### Microarrays

Experiments with phosphate-limited chemostats were performed as described in [16]. Extraction, labeling, and hybridization of RNA were performed as described in [16]. Microarrays were imaged using an Agilent SureScan G4900DA Microarray Scanner. Images were processed using Agilent Feature Extraction software and a custom R script. Fluorescence intensities were floored at a value of 50. The resulting data were hierarchically clustered (Pearson distance metric with average linkage) using Cluster 3.0 [89] and visualized in R using the *pheatmap* package.

### BAR-seq

To assess competitive fitness of strains in the YETI collections, pooled cultures were propagated at low density (OD_600_ < 0.05) in either SC +2% glucose +monosodium glutamate +ClonNAT or YNB +2% glucose +monosodium glutamate +ClonNAT either with or without addition of 100 nM β-estradiol, and > 2 × 10^6^ cells were collected at each time point. A wild type strain containing the Z_3_EV module integrated at the *HAP1* locus was spiked in (3 % of total cell number). For BAR-seq experiments performed in SC medium, samples were collected at t = 0, 9, 18, and 36 hours. In YNB medium, samples were collected at t = 0, 12, 24, and 48 hours. Genomic DNA was extracted using the YeaStar Genomic DNA kit (Zymo Research) and 5 ng of DNA were used to PCR amplify the barcode region with custom primers (1 min at 98°C; 20 cycles at 98°C for 30 s, 65°C for 30 s, 72°C for 45 s; 3 min at 72°C). Primers were designed as follows, where lowercase n indicates the position of the 8-bp index for multiplexing:

Forward:

5’-CAAGCAGAAGACGGCATACGAGATnnnnnnnnGTCTCGTGGGCTCGGAGATGTGTATAAGAGACAGCGGTGCGAGCGGATC-3’

Reverse:

5’-AATGATACGGCGACCACCGAGATCTACACnnnnnnnnTCGTCGGCAGCGTCAGATGTGTATAAGAGACAGGCACCAGGAACCATATA-3’

PCR products were purified with 2 volumes of RNAClean XP beads (Beckman Coulter) and then quality-checked and quantified on a Bioanalyzer (Agilent). Samples were pooled in equal proportions and purified further using the DNA Clean & Concentrator kit (Zymo Research). Barcodes were sequenced with a NovaSeq 6000 Sequencer (Illumina) according to manufacturer’s instructions and with addition of 30% PhiX Control v3 Library.

Samples were demuxed and barcodes were called with BARCAS [90] allowing up to two barcode mismatches. To generate heatmaps, the data were floored to the background level which was determined using the barcode count distribution of the time 0 samples. The barcode abundance for each strain was then normalized to the barcode abundance of the wild type strain and then normalized to the mean value of the three time 0 samples of the respective strain and displayed as a log2 fold change.

### Z_3_EV and Z_3_EB42 growth comparisons

All steps were carried out at 26°C. To test reversibility of essentiality, a panel of YETI-E diploids were pinned onto SC-URA+ClonNAT and incubated for two days. Colonies were then pinned onto sporulation medium, and incubated at for five days. To select for haploids containing Z_3_EV-controlled alleles, cells were pinned onto

SC-HIS-ARG-LYS-URA+canavanine+thialysine+ClonNAT medium and grown for two days in the presence of 10 nM β-estradiol. Colonies were then pinned to SC-HIS-ARG-LYS-URA+canavanine+thialysine+ClonNAT medium containing either 0 nM or 10 nM β-estradiol and grown for two days. These colonies were then again pinned onto the same medium containing either 0 nM or 10 nM β-estradiol. Plates were imaged after two days of growth. To test reversibility of Z_3_EB42-driven alleles, the same procedure was used, except for the addition of a higher concentration of β-estradiol (1 μM instead of 10 nM). Colonies on the perimeter of plates were removed from all analyses. The remaining colony sizes were quantified using *gitter [91].* We then computed the ratio of colony sizes in the presence and the absence of β-estradiol. Reversibility was defined using the following cutoffs:

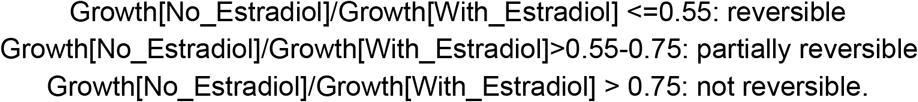

Identifying Z_3_EB42-driven alleles that had β-estradiol-dependent growth was done using the following procedure:

1. Colonies were pinned onto SC-URA+ClonNAT (as diploids) and grown at 26°C for two days
2. Colonies were pinned onto sporulation medium and grown at 26°C for five days to select for haploids with Z_3_EB42-driven alleles
3. Colonies were pinned on SC-HIS-ARG-LYS-URA+canavanine+thialysine+ClonNAT with 0, 10, 100, or 1000 nM β-estradiol and grown for two days
4. Images were taken and quantified using *gitter*.

### Gene Ontology

Gene Ontology (GO) enrichments were performed using GO-term Finder Version 0.86, which is made available through the *Saccharomyces cerevisiae* Genome Database (https://www.yeastgenome.org/goTermFinder).

### ROF1 Synthetic Dosage Lethality (SDL)

Strain BY6442 *MAT*α *HAP1+*::*natMX*::*ACT1*pr-Z_3_EV-*ENO2term ura3Δ0 can1*Δ::*STE2*pr-*SpHis5 his3Δ1 lyp1Δ URA3*-Z_3_pr*(6 binding sites)*-*ROF1* was derived from the YETI-NE heterozygote, ed2531, by tetrad dissection. SGA was done at 26°C using a standard protocol [92]: mating with the deletion mutant array (DMA) and the array of strains containing temperature sensitive alleles (TSA) on YPD (1 day), diploid selection on YPD+G418+ClonNAT (2 days), sporulation on spo + 1/4 G418 (5 days), first haploid selection on SD-his-ura-lys-arg+canavanine+thialysine (3 days), second haploid selection on SD-his-ura-lys-arg+canavanine+thialysine+G418 (2 days). For the final selection step, cells were pinned with 0.5 mm pins to SD-his-ura-lys-arg +canavanine+thialysine+G418+ClonNAT+0 nM β-estradiol and to SD-his-ura-lys-arg +canavanine+thialysine+G418+ClonNAT+1000 nM β-estradiol and incubated for 2 days (0 nM β-estradiol) and 5 days (1000 nM β-estradiol). Colony size was assessed using *gitter* [91] and SDL interactions were identified by dividing normalized colony size on plates with 1000 nM β-estradiol by normalized colony size on plates with 0 nM β-estradiol. Synthetic dosage lethal interactions were called for strains scoring < −0.08 in 2/2 replicate screens.

### Assessing expression of cells with different Z3 promoters driving mNeonGreen

Yeast strains with Z_3_EV and Z_3_EB42 transcriptional activators in combination with various Z_3_ promoters were grown to early log-phase in YNB medium, then diluted to OD_600_ = 0.05 in YNB containing a range of concentrations of β-estradiol. Cells were immediately inoculated in a 384-well imaging plate for growth at room temperature and imaged every hour with a Perkin Elmer Phenix automated confocal imaging system using digital phase contrast and EGFP channels. Cells were segmented in the digital phase contrast channel and average mNeonGreen intensity/cell was determined using the Harmony software package (Perkin-Elmer).

To assay the effects of β-estradiol removal, cells were grown to early log-phase in YNB containing 1000 nM β-estradiol, washed 3x in PBS, and inoculated in YNB lacking β-estradiol. Imaging and analysis were done as described above. Doubling time for cells in these conditions was 4-5 hours.

## Supporting information

p5820 map

HAP1 locus map

## Supplementary Figure Legends

**Figure S1: Characterization of Z_3_EV promoter behaviour**

**A.** Kinetics of GFP expression from the Z_3_EV promoter at 10 nM β-estradiol. Strains containing a Z_3_pr driving GFP (Y15292; blue) and control strain (Y15477; grey) were induced at time 0 with 10 nM β-estradiol in YNB medium. At each time point, cells were fixed and GFP intensity was measured by flow cytometry (AU = arbitrary units). Error bars represent standard deviation for three replicates. **B.** Characterization of Z_3_pr-*LEU2* strain. Tenfold serial dilutions of WT (Y15477) and the YETI-NE strain of *LEU2* are shown. The top row shows plates with no β-estradiol. Plates were incubated at 30°C for two days. **C.** Characterization Z_3_pr-*TPS2* strain. Y15477 and the YETI-NE strain of *TPS2* were spotted and grown at 30°C and 37°C as indicated on YPD with no β-estradiol or increasing concentrations of β-estradiol.

**Figure S2: Transcriptional induction of 18 YETI-NE strains**

**A.** Expression data of 18 strains during exponential growth in 1 mL batch culture of SC medium after 30 minutes of induction with varying concentrations of β-estradiol in a deep-well 96-well plate. The y-axis shows Transcripts Per Million (TPM) values from RNA-seq experiments and the x-axis shows concentrations of β-estradiol (nM). The dotted line indicates the endogenous level of expression for each gene in this experiment, calculated as the average TPM value from all strains in the panel that contain a native copy of the allele. **B.** Comparing Z_3_EV inducibility to endogenous gene expression. The graph shows maximum TPM values after β-estradiol induction for the 18 genes assayed in [A] (y-axis) versus native expression (x-axis). For genes with maximum endogenous expression level of <250 TPM, a linear model was fit (y = 87.96 + 4.79x, R^2^ = 0.76).

**Figure S3: Comparing growth and transcriptional induction for 18 YETI-NE strains**

Blue curves show growth data of 18 strains grown on minimal medium (on agar plates) with varying concentrations of β-estradiol. Error bars are +/- 1 standard deviation of the Area Under the Curve (AUGC) values acquired from eight independent colonies. Orange curves show log2 fold-change expression data of the same 18 strains from Figure S2A. Dotted vertical grey lines are shown at 0.1 nM β-estradiol in each subplot.

**Figure S4: Relationship between native gene expression and growth for YETI-E and other expression-perturbing collections of essential genes**

**A.** Box plots showing native levels of gene expression of genes from Lipson et al. (2009) (TPM, transcripts per million) as a function of the level of growth (from low to high, binned as growth deciles) of corresponding YETI-E strains at 0 nM β-estradiol. **B-E**. Box plots showing native levels of gene expression binned by growth characteristics for other yeast strain collections. **B.** The TET-allele collection. Genes not repressible with Tet-off (DOX non-responsive) and genes that are repressible with Tet-off (DOX-responsive) are shown. **C**. The DAmP allele collection. Box plots showing fitness of DAmP alleles as a function native expression level. **D.** TS-degron strains. Levels of native gene expression for genes for which the addition of an N-terminal degron does not affect growth versus those for which the degron confers temperature-sensitive growth are plotted. **E.** CRISPRi strains. Box plots showing fitness of CRISPRi alleles as a function native expression level.

**Figure S5: Z_3_pr-SHM2 is an adenine auxotroph**

**A.** Spot assays were performed with Z_3_pr-*MET6*, Z_3_pr-*TIP41*, and three independent colonies of Z_3_pr-*SHM2* on YNB and SC with increasing levels of β-estradiol. **B.** Spot assays were performed with Z_3_pr-*MET6*, Z_3_pr-*TIP41*, and four independent colonies of Z_3_pr-*SHM2* on SC, YNB, YNB with adenine, and YNB with glycine. **C.** Spot assays were performed with WT and four independent colonies of *shm2*Δ strains (two *MAT***a** and two MAT*α*) on SC, YNB, and YNB with adenine.

**Figure S6: Identification of genes that impair growth when overexpressed**

Growth (AUGC) in the presence (y-axis) and absence (x-axis) of β-estradiol for individual strains (points on the scatter plot). Blue points have a distance between −2,000 and 2,000 and are not considered dosage sensitive (Methods). Purple points have a distance less than −2,000 and yellow points have a distance greater than 2,000 and were identified as toxic strains. **A.** Toxic genes called for YETI-E strains grown at 100 nM and 0 nM. **B.** Toxic genes called for YETI-NE strains grown at 100 nM vs. 0 nM. **C.** Toxic genes called for YETI-E strains grown at β-estradiol concentrations (0.01, 0.03, 0.1, 0.3, 1, 3, 10, 30, 100, 300, 1000 nM) vs. 0 nM. **D.** Toxic genes called for YETI-NE strains grown at β-estradiol concentrations (1, 5, 10 and 100 nM vs. 0 nM).

**Figure S7: Non-essential toxic genes encode proteins that are bigger and more disordered than non-toxic genes**

**A.** Density plots of YETI-NE genes identified as non-toxic/toxic when overexpressed based on the percent of the protein that belongs to an intrinsically disordered region. **B.** Density plots of YETI-E genes identified as non-toxic/toxic when overexpressed based on the percent of the protein that belongs to an intrinsically disordered region. **C.** Density plots of YETI-NE genes identified as non-toxic/toxic when overexpressed based on protein length. **D.** Density plots of YETI-E genes identified as non-toxic/toxic when overexpressed based on protein length.

**Figure S8: Analysis of genes that impair growth when overexpressed**

**A.** Venn diagram of genes that are toxic upon overexpression from this study as compared to previous overexpression studies. **B.** Barplot of Z_3_EV toxic strains grouped by overlap with previous overexpression studies seen in **A**. For each grouping the percentage of Z_3_EV toxic strains that are toxic at only 100 nM β-estradiol (solid color) versus the percentage that are toxic at β-estradiol concentrations less than 100 nM (white) is shown.

**Figure S9: Scatterplots of log2-fold-changes of barcode frequencies in pooled culture (y-axis) after 24 hours of growth versus log2-fold-changes in total growth of strains on plates. The Spearman coefficient, and its significance, is shown in red.**

**Figure S10: Conditional growth of Z_3_EV-driven essential genes is not reversible in 16 of 24 strains tested**

**Figure S11: Reversibility analysis for 19 strains that impair growth when overexpressed**

**A.** The Response Ratio, *ResRatio_i_ = AUGC_i,0,0_ / AUGC_i,10,10_*, for each strain. For *AUGC*_i,j,k_, *i* is the gene, *j* is the concentration of β-estradiol during the first round of growth, and *k* is the level of β-estradiol during the second round of growth. For a given gene, the larger the *ResRatio* value, the greater the toxicity when the gene is induced. If log2(*ResRatio*) is negative, the gene improves growth when induced. Genes are ranked from least toxic to most toxic when induced in the presence of β-estradiol **B.** The Reversibility Ratio, *RevRatio_i_ = AUGC_i,10,0_ / AUGC_i,0,10i_*, for each strain. If log2(RevRatio_i_) > 0, the growth phenotype is at least partially reversible. **C-D.** Examples of reversible (C) and non-reversible (D) phenotypes. Barplots show total area under growth curve (AUGC) for cells grown at two β-estradiol concentrations and transferred to plates under four different conditions: 0 to 0_avg - a control transfer from no β-estradiol to no β-estradiol (no β-estradiol); 10 to 10_avg - another control transfer from β-estradiol to β-estradiol (continuous β-estradiol); 0 to 10_avg - transfer from no β-estradiol to β-estradiol (addition of β-estradiol) and; 10 to 0_avg - transfer from β-estradiol to no β-estradiol (removal of β-estradiol).

**Figure S12: Number of binding sites, URS1 presence, and ATF choice all affect mNeonGreen reporter induction time series**

Full time series collected for testing the relationship between ATF, URS1, and binding site copy number.

**Figure S13: Reversibility analysis of Z_3_EV and Z_3_EB42 using mNeonGreen reporter** Strains with all combinations of ATFs and promoters driving mNeonGreen were grown to early log phase in YNB + 1000 nM β-estradiol, washed 3x in PBS and resuspended in YNB without β-estradiol. Strains were imaged every hour using a Phenix automated confocal microscope and average GFP fluorescence/cell was calculated using Harmony software. Data are conditionally formatted such that high fluorescence is red and low fluorescence is blue. Numbers indicate arbitrary fluorescence units.

**Figure S14: Expression (TPM) levels of Core Essential Genes (CEG2)**

**Essential/Non-Essential genes in K562 cells.** The CEG2 essential list is from [94]. The median TPM of CEG2 Non-Essential genes is 5 TPM. The median TPM of CEG2 essential genes is 80 TPM.

## Tables

**Table S1: Summary of whole-genome sequencing data**

To identify possible aneuploidies and to confirm correct integration of the Z_3_EV promoter WGS was done on 107 strains, including some diploids and the haploids derived from them by SGA selection. If aneuploidy was observed (True in column 4), the aneuploid chromosome(s) is indicated in column 5.

**Table S2: List of β-estradiol-inducible strains**

**A.** Essential genes under the control of the Z_3_pr are available as **a/**α diploids.

**B, C.** Non-essential genes under the control of the Z_3_pr are available as **a**/α diploids (B) and as *MAT***a** haploids containing SGA markers (C). All essential and non-essential gene strains have a gene-specific barcode integrated upstream of the *URA3* selectable marker. Forward and reverse primers, as well as confirmation primers used in constructing the strains are shown. **D.** Strains whose construction was unsuccessful are shown. For 10 strains we constructed diploids but were not able to recover viable haploids following sporulation and SGA selection (last column).

**Table S3: Characterization of the YETI-E collection. Related to Figure 2.**

**A.** Area Under the Growth Curve values (AUGCs) for 975 YETI-E strains grown on SC medium with SGA selection. Each strain was assayed at 12 β-estradiol doses (0, 0.01, 0.03, 0.1, 0.3, 1, 3, 10, 30, 100, 300, 1000 nM). Following hierarchical clustering, each strain was assigned to one of five clusters. **B.** Normalized colony sizes for 54 strains from Cluster 3, which had shown little or no growth at all concentrations of β-estradiol that we attempted to remake. These strains have been removed from the clustergram in A. All reconstructed diploids were sporulated and SGA selection was used to select YETI-E haploids on 4 concentrations of β-estradiol (0, 10, 100, 1000 nM). Success at remaking was defined as AUGC >0.3 at any β-estradiol concentration. β-estradiol-dependent growth was defined as [maximum AUGC on any β-estradiol concentration] / [AUGC on 0 nM β-estradiol] > 2.

**Table S4: Comparison of inducible or repressible growth for essential genes with native gene expression levels for six expression-perturbing systems. Related to Figure 3 and Figure S4.**

Six independent studies assaying growth of strains with promoter-perturbing systems of essential genes are shown: YETI-E (yeast, β-estradiol-inducible) Tet-off (yeast, doxycycline-repressible), DAmP (reduces transcript abundance), TS-Degron (renders genes temperature-sensitive in *UBR1*-dependent manner), CRISPRi (yeast, repressible) and CRISPRi (*E. coli,* repressible). Transcript counts (TPM) for each gene expressed from its native promoter are shown. **A.** With the Z_3_EV system, strains in Essential Gene Clusters 1 and 2 were able to grow in the absence of β-estradiol, and strains in Clusters 4 and 5 were not able to grow in the absence of β-estradiol. **B.** With the Tet-off system [3], strains defined as Normal, Reverse and Slight were able to grow in the presence and absence of doxycycline. Strains defined as Severe and Very severe were unable to grow in the presence of doxycycline. **C.** DAmP (3’-UTR disruption) alleles. Median growth rates are from a competitive growth assay [57]. **D.** TS degron alleles. Phenotypes of strains with essential genes with degrons are from [7]. **E.** CRISPRi in yeast. Average gRNA depletion scores for essential genes are from [58]. **F.** In the CRISPRi study in *E. coli* [59], essential genes were defined using the Keio Collection. Based on a pooled CRISPRi experiment, these genes were classified as causing a dividing (Class D) or non-dividing phenotype (Class ND) when targeted by dCas9.

**Table S5: Auxotrophy in the YETI collection. Related to Figure 4 and Figure S5.**

**A.** AUGC values for Z-NE strains grown on YNB (0) and SC (1) medium with urea as nitrogen source at five β-estradiol concentrations (0, 1, 5, 10, 100 nM). **B.** Aux scores for YETI-NE strains. Column E indicates whether or not the corresponding deletion mutant was identified in the literature as an auxotroph (blue dots in Figure 4), not including genes in SGA selection pathways.

**Table S6: Overexpression toxicity in the YETI collection. Related to Figure S6, S7 and S8.**

**A.** Distance scores for YETI-E and YETI-NE strains grown on SC medium in the presence of 0 and 100 nM β-estradiol were determined as described in Materials and Methods. Genes with a distance score larger than 2,000 were considered toxic. Larger distance values represent toxic strains with a more extreme reduction in growth. **B.** GO enrichments were determined for the 987 YETI-E and YETI-NE genes with a toxic distance > 2000. Only GO terms with >2-fold enrichment and Bonferroni-corrected P-value <0.05 are shown. **C.** Percentage of each protein containing intrinsically disordered regions and length of protein (in amino acids) are shown. **D.** The 987 Z_3_EV toxic strains identified with 100 nM β-estradiol and their overlap with two other plasmid GAL promoter-based overexpression collections, GST and barFLEX, are shown. For each strain we compared growth in the presence of β-estradiol to growth in the absence of β-estradiol on synthetic complete medium; this is represented by the ‘distance’ column. The ‘overlap’ column indicates whether overexpression toxicity with Z_3_EV overlaps with overexpression toxicity in previous studies. Genes ‘Not screened in all 3 studies’ were not considered in our comparison. **E.** Distance scores at each concentration of β-estradiol were used to determine minimum toxic dose.

**Table S7: BAR-seq data for pooled YETI-NE collection at 0 and 100 nM β-estradiol. Related to Figure 5.**

Results for the YETI-NE collection grown in YNB and SC with 0 and 100 nM β-estradiol. **A.** Data [log2(fold-change) of barcode frequencies normalized to those at time 0] were hierarchically clustered to give the order of genes shown. **B.** Summary of properties of genes in each cluster. Properties of individual genes in each cluster are shown on subsequent sheets. GO enrichment of the genes in the 4 clusters is shown at the bottom. Only GO terms with >2-fold enrichment and Bonferroni-corrected P-value <0.05 are shown.

**Table S8: Microarray and SDL analysis of ROF1 overexpression. Related to Figure 6.**

**A.***ROF1* was induced with 1 μM β-estradiol and genome-wide transcriptional changes, normalized to time 0, were tracked over 90 minutes, identifying one regulon of Rof1-repressed genes (labeled R). **B.** GO enrichment of the genes in the Rof1-repressed regulon is shown. Only GO terms with >2-fold enrichment and Bonferroni-corrected P-value <0.05 are shown. **C.** SDL interactions with *ROF1*. Scores for two replicates with associated P-values are shown. Only strains with scores below our intermediate cutoff (−0.08) in 2/2 reps are shown. Array indicates the array on which the interaction was identified. TSA: temperature-sensitive array; DMA: deletion mutant array. Some mutants are present on both arrays. Biological role column shows only genes in the CWI (cell wall integrity/PKC) pathway and genes involved in chromatin/general transcription. Protein complex information from SGD is shown for genes with roles in chromatin/general transcription.

**Table S9: Reversibility of Z_3_EV system for growth of strains with essential genes. Related to Figure S10.** Reversibility of growth for essential genes was tested by transferring 24 YETI-E strains that initially had β-estradiol-dependent growth from plates with 10 nM to plates with 0 nM β-estradiol. Reversibility was quantified as follows using colony size on 0 nM and 10 nM β-estradiol: Ratio 0 nM /10 nM <=0.55: reversible; ratio 0 nM /10 nM >0.55-0.75: partially reversible; ratio 0 nM /10 nM>0.75: not reversible.

**Table S10: Z_3_EB42 β-estradiol-dependent growth and reversibility. Related to Figure 8.**

One hundred and thirteen **a**/α diploid strains were constructed with essential genes under the control of the Z_3_EB42 system. Diploids were sporulated and SGA selection was used to select haploid Z_3_EB42 strains on 0 nM and 1000 nM β-estradiol. Reversibility of growth for essential genes was tested by transferring 48 Z_3_EB42 strains that initially had β-estradiol-dependent growth from plates with 1000 nM to 0 nM β-estradiol. Initial β-estradiol-dependence/ reversibility were quantified as follows using colony size on 0 nM and 1000 nM β-estradiol: Ratio 0 nM / 1000 nM <=0.55: β-estradiol-dependent / reversible; ratio 0 nM / 1000 nM >0.55-0.75: partially β-estradiol-dependent / partially reversible; ratio 0 nM / 1000nM >0.75: constitutive / not reversible. We further labelled strains as slow growth (either β-estradiol-dependent or constitutive) for normalized colony size in 1000 nM β-estradiol <0.7.

## Files

File 1: URA3-Z_3_pr sequence

File 2: HAP1 locus sequence

## Data accession

Next-generation sequencing and microarray data can be found at the Gene Expression Omnibus with accession number GSE158319: https://www.ncbi.nlm.nih.gov/geo/query/acc.cgi?acc=GSE158319

## Acknowledgements

The authors acknowledge Sean Hackett, Brian Feng, Chiraj Dalal, and Meng Jin for providing critical feedback on the manuscript, and Bernd Wranik for providing assistance with the *ROF1* experiment. We thank Andrew Bognar for discussions about *SHM2*, Jef Boeke for information and reagents associated with URS1, and Adam Baker for his design insights and contributions to figure preparation.

---

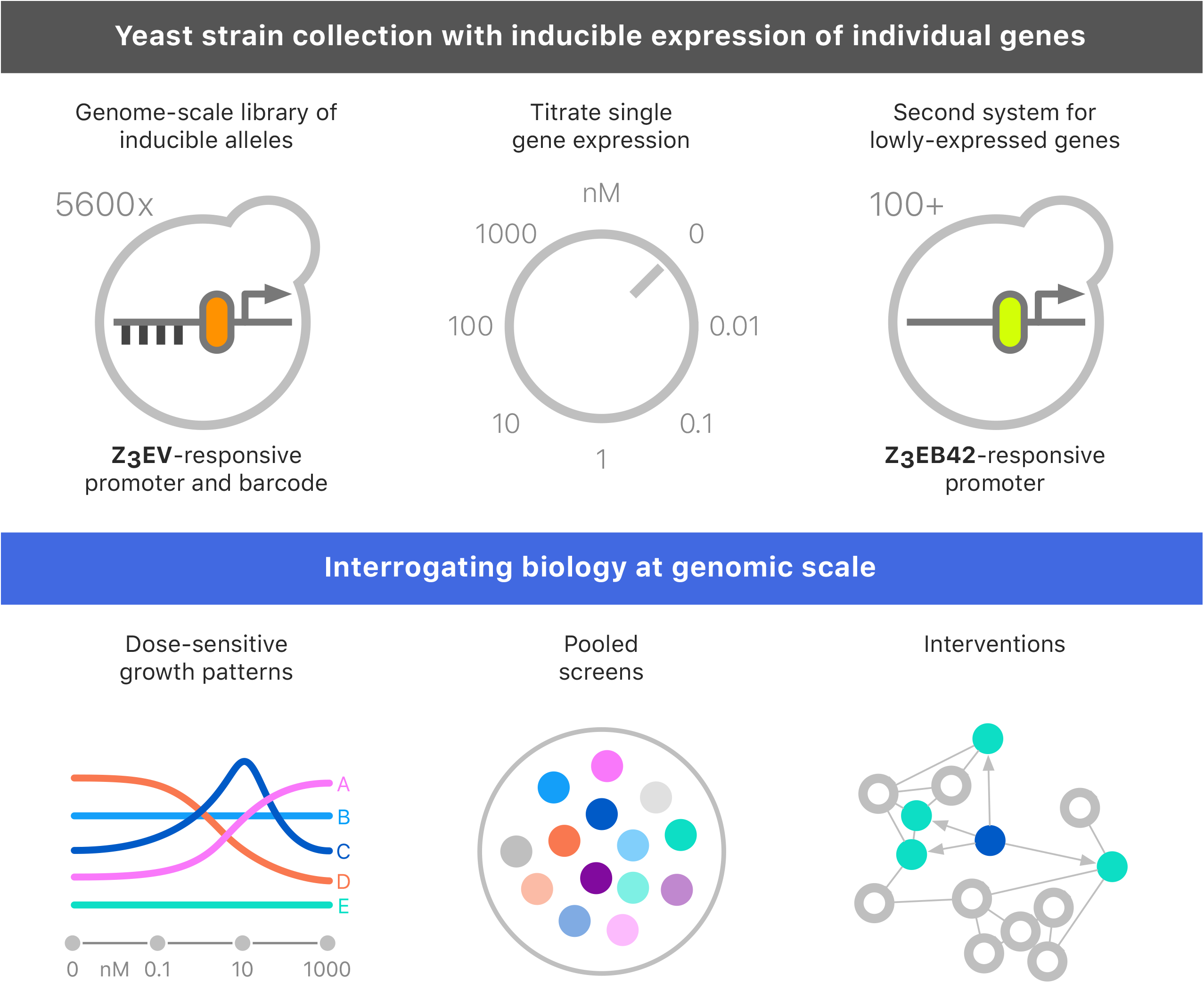

