## Supplementary material for "A genome-scale yeast library with inducible expression of individual genes": Figure S8

**a** Overlap of genes that impair growth when overexpressed

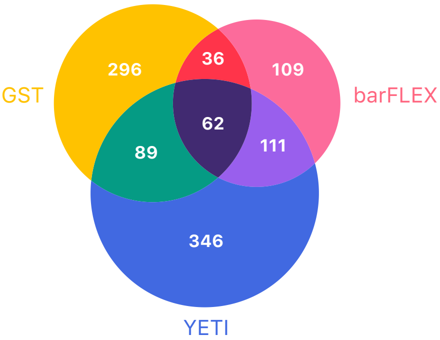


**b** Percentage of YETI strains toxic at or under 100 nM ß-estradiol

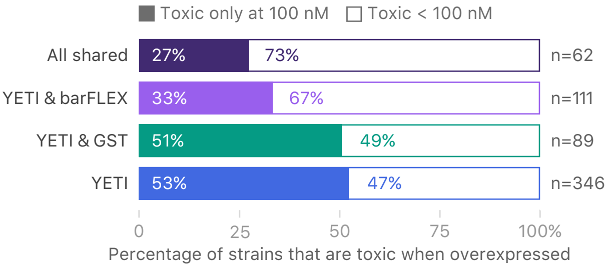
