## Supplemental Text for "A genome-scale yeast library with inducible expression of individual genes"

### Sequence information

#### p5820, $Z_3$ promoter sequence (6 binding sites)

The sequence of $Z_3$ (6 binding sites) promoter is shown below. Modified $UAS_{GAL}$ for $Z_3$EV binding is highlighted in green. XbaI and NotI restriction sites are colored with gray. A Gal4 binding site outside of XbaI and NotI restriction sites was also removed. As for 2 and 4 binding sites, we removed appropriate number of binding sites from 6 binding sites.

TTATATTGAATTTTCAAAAATTCTTACTTTTTTTTTGGATGGACGCAAAGAAGTTTAATAATCATATTACATGGCATTACCACCATAT
ACATATCCATATACATATCCATATCTAATCTTACTTATATGTTGTGGAAATGTAAAGAGCCCCATTATCTTAGCCTAAAAAAACCTT
CTCTTTGGAACTTTCAGTAATACGCTTAACTGCTCATTGCTATATTGAAGTGCGGCCGCGTGGGCGTGCGTGGGCGGGCGTGG
GCGTGCGTGGGCGGGCGTGGGCGTGCGTGGGCGTCTAGACCGTGCGTCCTCGTCTTCACCGGTCGCGTTCCTGAAACGCAG
ATGTGCCTCGCGCCGCACTGCTAGCAACAATAAAGATTCTACAATACTAGCTTTTATGGTTATGAAGAGGAAAAATTGGCAGTAA
CCTGGCCCCACAAACCTTCAAATTAACGAATCAAATTAACAACCATAGGATGATAATGCGATTAGTTTTTTAGCCTTATTTCTGGG
GTAATTAATCAGCGAAGCGATGATTTTTGATCTATTAACAGATATATAAATGGAAAAGCTGCATAACCACTTTAACTAATACTTTCA
ACATTTTCAGTTTGTATTACTTCTTATTCAAATGTCATAAAAGTATCAACAAAAAAATTGTTAATATACCTCTATACTTTAACGTCAAG
GAGAAAAAACTATA

#### p7418, $Z_3$ promoter sequence (2 binding sites)

TTATATTGAATTTTCAAAAATTCTTACTTTTTTTTTGGATGGACGCAAAGAAGTTTAATAATCATATTACATGGCATTACCACCATAT
ACATATCCATATACATATCCATATCTAATCTTACTTATATGTTGTGGAAATGTAAAGAGCCCCATTATCTTAGCCTAAAAAAACCTT
CTCTTTGGAACTTTCAGTAATACGCTTAACTGCTCATTGCTATATTGAAGTGCGGCCGCGTGGGCGTGCGTGGGCGGTCTAGAC
CGTGCGTCCTCGTCTTCACCGGTCGCGTTCCTGAAACGCAGATGTGCCTCGCGCCGCACTGCTAGCAACAATAAAGATTCTAC
AATACTAGCTTTTATGGTTATGAAGAGGAAAAATTGGCAGTAACCTGGCCCCACAAACCTTCAAATTAACGAATCAAATTAACAA
CCATAGGATGATAATGCGATTAGTTTTTTAGCCTTATTTCTGGGGTAATTAATCAGCGAAGCGATGATTTTTGATCTATTAACAGA
TATATAAATGGAAAAGCTGCATAACCACTTTAACTAATACTTTCAACATTTTCAGTTTGTATTACTTCTTATTCAAATGTCATAAAAG
TATCAACAAAAAAATTGTTAATATACCTCTATACTTTAACGTCAAGGAGAAAAAACTATA

#### p7460, $Z_3$ (6 binding sites)-URS1 promoter sequence

Modified $UAS_{GAL}$ forATF binding is highlighted in green. XbaI and NotI restriction sites are colored with gray. URS1 sequence of CAR1 is highlighted in blue.

TTATATTGAATTTTCAAAAATTCTTACTTTTTTTTTGGATGGACGCAAAGAAGTTTAATAATCATATTACATGGCATTACCACCATAT
ACATATCCATATACATATCCATATCTAATCTTACTTATATGTTGTGGAAATGTAAAGAGCCCCATTATCTTAGCCTAAAAAAACCTT
CTCTTTGGAACTTTCAGTAATACGCTTAACTGCTCATTGCTATATTGAAGTGCGGCCGCGTGGGCGTGCGTGGGCGGGCGTGG
GCGTGCGTGGGCGGGCGTGGGCGTGCGTGGGCGTCTAGACCGTGCGTCCTCGTCTTCACCGGTCGCGTTCCTGAAACGCAG
ATGTGCCTCGCGCCGCACTGCTAGCAACAATAAAGATTCTACAATACTAGCTTTTATGGTTATGAAGAGGAAAAATTGGCAGTAA
CCTGGCCCCACAAACCTTCAAATTAACGAATCAAATTAACAACCATAGGATGATAATGCGATTAGTTTTTTAGCCTTATTTCTGGG
GTAATTAATCAGCGAAGCGATGATTTTTGATCTATTAACAGATATATAAATGGAAAAGCTGCATAACCACTTTAACTAATACTTTCA
ACATTTTCAGTTTGTATTACTTCTTATTCAAATGTCATAAAAGTATCAACAAAAAAATTGTTAATATACCTCTATATCAACAGCGCATC
GCCGCTCGCTGAATTTTTCACTTAGCGGTAGCCGCCGAGGGGTCTAAAGAGTATATAAGCAGAGCTTGCGGCCCACTTTCTATC
AAGATCTAAGACTGTTTCTCTTCTCTTGGTCTGTATATGTTTTCTCAAAGTTAGCAGAAACAACAACAACAACTATATCAATAACAA
TAACTACTATCAAGCTTTAACGTCAAGGAGAAAAAACTATA

#### p7288, $Z_3$ (2 binding sites)-URS1 promoter sequence

Modified $UAS_{GAL}$ for ATF binding is highlighted in green. XbaI and NotI restriction sites are colored with gray. URS1 sequence of CAR1 is highlighted in blue.

TTATATTGAATTTTCAAAAATTCTTACTTTTTTTTTGGATGGACGCAAAGAAGTTTAATAATCATATTACATGGCATTACCACCATAT
ACATATCCATATACATATCCATATCTAATCTTACTTATATGTTGTGGAAATGTAAAGAGCCCCATTATCTTAGCCTAAAAAAACCTT
CTCTTTGGAACTTTCAGTAATACGCTTAACTGCTCATTGCTATATTGAAGTGCGGCCGCGTGGGCGTGCGTGGGCGGTCTAGAC
CGTGCGTCCTCGTCTTCACCGGTCGCGTTCCTGAAACGCAGATGTGCCTCGCGCCGCACTGCTAGCAACAATAAAGATTCTAC

AATACTAGCTTTTATGGTTATGAAGAGGAAAAATTGGCAGTAACCTGGCCCCACAAACCTTCAAATTAACGAATCAAATTAACAA
CCATAGGATGATAATGCGATTAGTTTTTTAGCCTTATTTCTGGGGTAATTAATCAGCGAAGCGATGATTTTTGATCTATTAACAGA
TATATAAATGGAAAAGCTGCATAACCACTTTAACTAATACTTTCAACATTTTCAGTTTGTATTACTTCTTATTCAAATGTCATAAAAG
TATCAACAAAAAATTGTTAATATACCTCTATATCAACAGCGCATCGCCGCTCGCTGAATTTTTCACTTAGCGGTAGCCGCCGAGG
GGTCTAAAGAGTATATAAGCAGAGCTTGCGGCCCACTTTCTATCAAGATCTAAGACTGTTTCTCTTCTCTTGGTCTGTATATGTTT
TCTCAAAGTTAGCAGAAACAACAACAACAACTATATCAATAACAATAACTACTATCAAGCTTTAACGTCAAGGAGAAAAAACTATA

#### $Z_3$EB42 sequence

The *ACT1* promoter is shown in blue; the $Z_3$ DNA binding domain is shown in green; the human Estrogen Receptor is shown in magenta; the B42 activation domain is in red; the *ENO2* terminator is in orange. The transcription factor was integrated 300 bp downstream of functional *HAP1*. 300 bp of downstream sequence of *HAP1* was also added downstream of the $Z_3$EB42 transcription factor.

GCCTCTACCTTGCAGACCCATATAATATAATAACTAAATAAGTAAATAAGACACACGCGAGAACATATATACACAATTACAGTAAC
AATAACAAGAGGACAGATACTACCAAAATGTGTGGGGAAGCGGGTAAGCTGCCACAGCAATTAATGCACAACATTTAACCTACA
TTCTTCCTTATCGGATCCTCAAAACCCTTAAAAACATATGCCTCACCCTAACATATTTTCCAATTAACCCTCAATATTTCTCTGTCA
CCCGGCCTCTATTTTCCATTTTCTTCTTTACCCGCCACGCGTTTTTTTCTTTCAAATTTTTTTCTTCCTTCTTCTTTTTCTTCCACGT
CCTCTTGCATAAATAAATAAACCGTTTTGAAACCAAACTCGCCTCTCTCTCTCCTTTTTGAAATATTTTTGGGTTTGTTTGATCCTT
TCCTTCCCAATCTCTCTTGTTTAATATATATTCATTTATATCACGCTCTCTTTTTATCTTCCTTTTTTTCCTCTCTCTTGTATTCTTCC
TTCCCCTTTCTACTCAAACCAAGAAGAAAAAGAAAAGGTCAATCTTTGTTAAAGAATAGGATCTTCTACTACATCAGCTTTTAGAT
TTTTCACGCTTACTGCTTTTTTCTTCCCAAGATCGAAAATTTACTGAATTAACAGGGCCCCCCCTCGAGGTCGACGGTATCGATA
AGCTTGAAGCAAGCCTCCTGAAAGATGGGTACCCGCCCATATGCTTGCCCTGTCGAGTCCTGCGATCGCCGCTTTTCTCGCTC
GGATGAGCTTACCCGCCATATCCGCATCCATACCGGTCAGAAGCCCTTCCAGTGTCGAATCTGCATGCGTAACTTCAGTCGTAG
TGACCACCTTACCACCCACATCCGCACCCACACAGGCGAGAAGCCTTTTGCCTGTGACATTTGTGGGAGGAAGTTTGCCAGGA
GTGATGAACGCAAGAGGCATACCAAAATCCATACAGGTGGCGGAGGCACACCTGCAGCTGCGTCGACTCTAGAGGATCCATCT
GCTGGAGACATGAGAGCTGCCAACCTTTGGCCAAGCCCGCTCATGATCAAACGCTCTAAGAAGAACAGCCTGGCCTTGTCCCT
GACGGCCGACCAGATGGTCAGTGCCTTGTTGGATGCTGAGCCCCCCATACTCTATTCCGAGTATGATCCTACCAGACCCTTCA
GTGAAGCTTCGATGATGGGCTTACTGACCAACCTGGCAGACAGGGAGCTGGTTCACATGATCAACTGGGCGAAGAGGGTGCC
AGGCTTTGTGGATTTGACCCTCCATGATCAGGTCCACCTTCTAGAATGTGCCTGGCTAGAGATCCTGATGATTGGTCTCGTCTG
GCGCTCCATGGAGCACCCAGTGAAGCTACTGTTTGCTCCTAACTTGCTCTTGGACAGGAACCAGGGAAAATGTGTAGAGGGCA
TGGTGGAGATCTTCGACATGCTGCTGGCTACATCATCTCGGTTCCGCATGATGAATCTGCAGGGAGAGGAGTTTGTGTGCCTCA
AATCTATTATTTTGCTTAATTCTGGAGTGTACACATTTCTGTCCAGCACCCTGAAGTCTCTGGAAGAGAAGGACCATATCCACCG
AGTCCTGGACAAGATCACAGACACTTTGATCCACCTGATGGCCAAGGCAGGCCTGACCCTGCAGCAGCAGCACCAGCGGCTG
GCCCAGCTCCTCCTCATCCTCTCCCACATCAGGCACATGAGTAACAAAGGCATGGAGCATCTGTACAGCATGAAGTGCAAGAA
CGTGGTGCCCCTCTATGACCTGCTGCTGGAGATGCTGGACGCCCACCGCCTACATGCGCCCACTAGCCGTGGAGGGGCATCC
GTGGAGGAGACGGACCAAAGCCACTTGGCCACTGCGGGCTCTACTTCATCGGGTATCAATAAAGAcATCGAGGAGTGCAATGC
CATCATTGAGCAGTTTATCGACTACCTGCGCACCGGACAGGAGATGCCGATGGAAATGGCGGATCAGGCGATTAACGTGGTGC
CGGGCATGACGCCGAAAACCATTCTTCACGCCGGGCCGCCGATCCAGCCTGACTGGCTGAAATCGAATGGTTTTCATGAAATT
GAAGCGGATGTTAACGATACCAGCCTCTTGCTGAGTGGAGATGCCTCCAAGCTTTAAATCCCCGCGTGCTTGGCCGGCCGTAG
TGCTTTTAACTAAGAATTATTAGTCTTTTCTGCTTATTTTTTCATCATAGTTTAGAACACTTTATATTAACGAATAGTTTATGAATCT
ATTTAGGTTTAAAAATTGATACAGTTTTATAAGTTACTTTTTCAAAGACTCGTGCTGTCTATTGCATAATGCACTGGAAGGGGAAA
AAAAAGGTGCACACGCGTGGCTTTTTCTTGAATTTGCAGTTTGAAAAATAACTACATGGATGATAAGAAAACATGGAGTACAGTC
ACTTTGAGAACCTTCAATCAGCTGGTAACGTCTTCGTTAATTGGATACTCAAAAAAGATGGATAGCATGAATCACAAGATGGAAG
GAAATGCGGGCCACGACCACAGTGATATGCATATGGGAGATGGAGATGATACCTGTTCGATGAATATGCTATTTTCGTGGTCAT
ACAAGAATACGTGTGTCGTCTTTGAATGGTGGCATATCAAGACCCTGCCTGGACTGAT

### Whole-genome sequencing of strains

Additional QC of the library was done by whole-genome sequencing of a subset of strains to check for correct insertion of $Z_3$EVpr as well as to look for any potential signs of aneuploidy. A total of 107 strains were selected to cover both essential and non-essential strains: 27 essential diploid strains and 40 non-essential strains in diploid and haploid. Overall, 95.3% of strains show proper insertion and attachment of the $Z_3$EVpr. 86.0% of all strains in our panel showed no signs of aneuploidy. Strains in this panel are enriched for haploids involved in pathways that get selected against when undergoing the SGA selection procedure (URA, HIS, LYS, and ARG selections/genes). If we remove those strains from our analysis, then 95.9% of all strains tested showed no signs of aneuploidy in our panel.

### Media recipes

**Z3-E and Z3-NE Diploid Maintenance Medium**

YPD+ClonNAT

For 1L

- In 900 mL water, add
  10 g Yeast Extract
  20 g Bacto Peptone
  20 g Agar
- autoclave
- Add 100 mL 20% glucose (w/v)
- Mix well and cool to ~60°C
- Add 1 mL 1000X ClonNAT (100mg/mL; final concentration 100 μg/mL)

**Sporulation Medium**

For 1L

- 10 g potassium acetate
- 1 g yeast extract
- 20 g bacto agar to 1L water
- After autoclaving, cool to 50°C
- 2.5 mL of 20% glucose (w/v)
- 1.67 mL 100X histidine (8.56 g/L; final concentration ~14 mg/L)
- 166.7 μL of 1000X G418 stock solution (300 mg/mL; final concentration 50 μg/L)

**Z3-E Haploid Selection Medium**

**(SC-arg-his-lys-ura)+YNB+glucose+msg+canavanine+thialysine+ClonNAT+β-estradiol**

For 5L

- In 3L water, add
  8.5g YNB w/o Amino Acids w/o Ammonium Sulfate
  100g agar
  milliQ water up to 3.5L
- Autoclave, cool to 50°C, and add
  500mL 10X SC-arg-his-lys-ura (manufacturer's recommendation)
  500mL 20% glucose (w/v)
  500mL 10X msg (760 mM; final concentration 76mM)
  5mL 1000X canavanine (60 mg/mL; final concentration 60 μg/mL)
  2.5mL 2000X thialysine (100 mg/mL; final concentration 100 μg/mL)
  5mL 1000X ClonNAT (100mg/mL; final concentration 100 μg/mL)
  5mL 1000X β-estradiol (1000X of final concentration assayed, in ethanol)

**Z3-NE Haploid Maintenance Medium**

**SC+YNB+glucose+msg+ClonNAT**

For 5L

- In 3L water, add
  8.5g YNB w/o Amino Acids w/o Ammonium Sulfate
  100g agar
  milliQ water up to 3.5L
- Autoclave, cool to 50°C, and add
  500mL 10X SC (manufacturer's recommendation)

500mL 20% glucose (w/v)
500mL 10X msg (760 mM; final concentration 76mM)
5mL 1000X ClonNAT (100mg/mL; final concentration 100 μg/mL)

**Z3-NE Haploid SC Medium**
**SC+YNB+glucose+msg+ClonNAT+β-estradiol**
For 5L
- In 3L water, add
  8.5g YNB w/o Amino Acids w/o Ammonium Sulfate
  100g agar
  milliQ water up to 3.5L
- Autoclave, cool to 50°C, and add
  500mL 10X SC (manufacturer's recommendation)
  500mL 20% glucose (w/v)
  500mL 10X msg (760 mM; final concentration 76mM)
  5mL 1000X ClonNAT (100mg/mL; final concentration 100 μg/mL)
  5mL 1000X β-estradiol (1000X of final concentration assayed, in ethanol)

**Z3-NE Haploid Minimal Medium**
**YNB+glucose+msg+ClonNAT+β-estradiol**
For 5L
- In 3.5L water, add
  8.5g YNB w/o Amino Acids w/o Ammonium Sulfate
  100g agar
  milliQ water up to 4L
- Autoclave, cool to 50°C, and add
  500mL 20% glucose (w/v)
  500mL 10X msg (760 mM; final concentration 76mM)
  5mL 1000X ClonNAT (100mg/mL; final concentration 100 μg/mL)
  5mL 1000X β-estradiol (1000X of final concentration assayed, in ethanol)

**10X SC solution (for preparation of 1L)**
- Add 20 g of SC mix (Sunrise Cat# 1300-030) into a beaker with stir bar
- Add 900 mL of water
- Stir until dissolved (heat liquid up to 60°C to dissolve)
- Transfer into a graduated cylinder, and add water to 1000 mL
- Filter sterilize using 1000 mL stericup

**10X SC-arg-his-lys-ura solution (for preparation of 1L)**
- Add 16.6 g of SC-arg-his-lys-ura mix (Sunrise Cat# 6103-030) into a beaker with stir bar
- Add 900 mL of water
- Stir until dissolved (heat liquid up to 60°C to dissolve)
- Transfer into a graduated cylinder, and add water to 1000 mL
- Filter sterilize using 1000 mL stericup

### Yeast Strains

| Strain | Genotype | Source |
|---|---|---|
| RCY1972 | *MAT***a** *his3Δ1 HAP1+* | Gift from Amy Caudy |
| BY6442 | *MATα URA3::$Z_3$pr-ROF1 HAP1+::natMX::pACT1-$Z_3$EV-ENO2term ura3Δ0 can1Δ::STE2pr-Sphis5 his3Δ1 lyp1Δ0* | this study |
| DBY12394 | *MAT**a** ura3Δ0 leu2Δ0 pr-ACT1-$Z_3$EV-NatMX* | McIsaac et al. (2013) |
| Y7092 | *MAT***a** *can1Δ::STE2pr-Sphis5 lyp1Δ his3Δ1 leu2Δ0 ura3Δ0 met15Δ0* | Costanzo et al. (2010) |
| Y14851 | *MAT***a** *HAP1+::natMX::ACT1*pr-$Z_3$EV-*ENO2term ura3Δ0 can1ΔSTE2*pr-*Sphis5 his3Δ1 lyp1Δ* | this study |
| Y14537 | *MATα ura3Δ0 his3Δ1 HAP1+* | this study |
| Y14789 | *MAT***a***/α HAP1+::natMX::ACT1*pr-$Z_3$EV-*ENO2term/HAP1+ ura3Δ0/ura3Δ0 can1Δ::STE2*pr-*Sphis5/CAN1+ his3Δ1/ his3Δ1 lyp1Δ/LYP1+* | this study |
| Y15090 | *MAT***a***/α HAP1-natMX-ACT1pr-Z3EV-ENO2term/HAP1 ura3Δ0/URA3 can1Δ::STE2pr-Sphis5/CAN1 his3Δ1/his3Δ1 lyp1Δ/LYP1* | this study |
| Y15292 | *MAT***a** *HAP1+::natMX::ACT1*pr-$Z_3$EV-*ENO2term ura3Δ0 can1ΔSTE2*pr-*Sphis5 his3Δ1 lyp1Δ hoΔ::URA3::$Z_3$pr(6 binding sites)-GFP* | this study |
| Y15260 | *MAT***a***/α HAP1+::natMX-pACT1-$Z_3$EB42ENO2term /HAP1+ his3Δ1/his3Δ1 ura3Δ0/ura3Δ0 lyp1Δ/LYP1 can1ΔSte2pr-Sphis5/CAN1 leu2Δ/leu2Δ URA3-$Z_3$pr(2 binding sites)-CAR1URS1-PBR1* | this study |
| Y15474 | *MAT***a***/α HAP1+::natMX-pACT1-$Z_3$EB42-ENO2term /HAP1+ his3Δ1/his3Δ1 ura3Δ0/ura3Δ0 lyp1Δ/LYP1 can1ΔSte2pr-Sphis5/CAN1 leu2Δ/leu2Δ URA3-$Z_3$pr(2 binding sites)-CAR1URS1-RAD53* | this study |
| Y15390 | *MAT***a***/α HAP1+::natMX-pACT1-$Z_3$EB42-ENO2term /HAP1+ his3Δ1/his3Δ1 ura3Δ0/ura3Δ0 lyp1Δ/LYP1 can1ΔSte2pr-Sphis5/CAN1 leu2Δ/leu2Δ URA3-$Z_3$pr(2 binding sites)-CAR1URS1-CIA2* | this study |
| Y15475 | *MAT***a***/α HAP1+::natMX-pACT1-$Z_3$EB42-ENO2term /HAP1+ his3Δ1/his3Δ1 ura3Δ0/ura3Δ0 lyp1Δ/LYP1 can1ΔSte2pr-Sphis5/CAN1 leu2Δ/leu2Δ URA3-$Z_3$pr(2 binding sites)-CAR1URS1-IPL1* | this study |
| Y15476 | *MAT***a***/α HAP1+::natMX-pACT1-$Z_3$EB42-ENO2term /HAP1+ his3Δ1/his3Δ1 ura3Δ0/ura3Δ0 lyp1Δ/LYP1 can1ΔSte2pr-Sphis5/CAN1 leu2Δ/leu2Δ URA3-$Z_3$pr(2 binding sites)-CAR1URS1-HYP2* | this study |
| Y15477 | *MAT***a** *HAP1+::natMX::ACT1*pr-*$Z_3$EV-ENO2term can1ΔSTE2*pr-*Sphis5 his3Δ1 lyp1Δ URA3+* | this study |
| Y15478 | *MAT***a***/α HAP1+::natMX::ACT1pr-$Z_3$EV-ENO2term/HAP1+ ura3Δ0/ura3Δ0 can1ΔSTE2pr-Sphis5/CAN1+ his3Δ1/ his3Δ1 lyp1Δ/LYP1+ URA3-$Z_3$pr(2 binding sites)-RAD53* | this study |
| Y15479 | *MAT***a***/α HAP1+::natMX::ACT1pr-$Z_3$EV-ENO2term/HAP1+ ura3Δ0/ura3Δ0 can1ΔSTE2pr-Sphis5/CAN1+ his3Δ1/ his3Δ1 lyp1Δ/LYP1+ URA3-$Z_3$pr(4 binding sites)-RAD53* | this study |

| | | |
|---|---|---|
| cDBY0776 | *MAT***a** *HAP1+ shm2Δ::kanMX* | this study |
| cDBY0777 | *MAT***a** *HAP1+ shm2Δ::kanMX* | this study |
| cDBY0778 | *MATα HAP1+ shm2Δ::kanMX* | this study |
| cDBY0779 | MAT*α HAP1+ shm2Δ::kanMX* | this study |
| Y15513 | *MAT***a** *HAP1+*::*natMX*::*ACT1*pr-$Z_3$EV-*ENO2term ura3Δ0 can1ΔSTE2*pr-*Sphis5 his3Δ1 lyp1Δ hoΔ*::*URA3*-$Z_3$pr*(6 binding sites)*-mNeonGreen | this study |
| Y15515 | *MAT***a** *HAP1+*::*natMX*::*ACT1*pr-$Z_3$EV-*ENO2term ura3Δ0 can1ΔSTE2*pr-*Sphis5 his3Δ1 lyp1Δ hoΔ*::*URA3*-$Z_3$pr*(2 binding sites)*-mNeonGreen | this study |
| Y15516 | *MAT***a** *HAP1+*::*natMX*::*ACT1*pr-$Z_3$EV-*ENO2term ura3Δ0 can1ΔSTE2*pr-*Sphis5 his3Δ1 lyp1Δ hoΔ*::*URA3*-$Z_3$pr*(6 binding sites-CAR1URS1)*-mNeonGreen | this study |
| Y15517 | *MAT***a** *HAP1+*::*natMX*::*ACT1*pr-$Z_3$EV-*ENO2term ura3Δ0 can1ΔSTE2*pr-*Sphis5 his3Δ1 lyp1Δ* hoΔ::*URA3*-$Z_3$pr*(2 binding sites-CAR1URS1)*-mNeonGreen | this study |
| Y15518 | *MAT***a** *HAP1+*::*natMX*::*ACT1*pr-$Z_3$EB42-*ENO2term ura3Δ0 can1ΔSTE2*pr-*Sphis5 his3Δ1 lyp1Δ hoΔ*::*URA3*-$Z_3$pr*(6 binding sites)*-mNeonGreen | this study |
| Y15520 | *MAT***a** *HAP1+*::*natMX*::*ACT1*pr-$Z_3$EB42-*ENO2term ura3Δ0 can1ΔSTE2*pr-*Sphis5 his3Δ1 lyp1Δ hoΔ*::*URA3*-$Z_3$pr*(2 binding sites)*-mNeonGreen | this study |
| Y15521 | *MAT***a** *HAP1+*::*natMX*::*ACT1*pr-$Z_3$EB42-*ENO2term ura3Δ0 can1ΔSTE2*pr-*Sphis5 his3Δ1 lyp1Δ hoΔ*::*URA3*-$Z_3$pr*(6 binding sites-CAR1URS1)*-mNeonGreen | this study |
| Y15522 | *MAT***a** *HAP1+*::*natMX*::*ACT1*pr-$Z_3$EB42-*ENO2term ura3Δ0 can1ΔSTE2*pr-*Sphis5 his3Δ1 lyp1Δ hoΔ*::*URA3*-$Z_3$pr*(2 binding sites-CAR1URS1)*-mNeonGreen | this study |

### Plasmids

| Plasmid | Description | Source |
|---|---|---|
| pRB3460 | Original promoter that is inducible by $Z_3$EV ($Z_3$pr); contains *kanMX* | McIsaac et al. (2013) |
| p5820 | Template for PCR targeting; contains *URA3* upstream of $Z_3$pr | this study |
| p7418 | URA3-$Z_3$pr (2 binding sites) | this study |
| p7288 | URA3-$Z_3$pr (CAR1 URS + 2 Zif268 Binding Sites) | this study |
| p7460 | URA3-$Z_3$pr (CAR1 URS + 6 Zif268 Binding Sites) | this study |
